# Non-parametric mixture models identify trajectories of childhood immune development relevant to asthma and allergy

**DOI:** 10.1101/237073

**Authors:** Howard H.F. Tang, Shu Mei Teo, Danielle C.M. Belgrave, Michael D. Evans, Daniel J. Jackson, Marta Brozynska, Merci M.H. Kusel, Sebastian L. Johnston, James E. Gern, Robert F. Lemanske, Angela Simpson, Adnan Custovic, Peter D. Sly, Patrick G. Holt, Kathryn E. Holt, Michael Inouye

**Affiliations:** School of BioSciences, The University of Melbourne, Parkville, Victoria, Australia; Baker Heart and Diabetes Institute, Melbourne, Victoria, Australia; Bio21 Molecular Science and Biotechnology Institute, The University of Melbourne, Parkville, Victoria, Australia; Department of Public Health and Primary Care, University of Cambridge, Cambridge, United Kingdom; Department of Paediatrics, Imperial College, London, United Kingdom; University of Wisconsin School of Medicine and Public Health, Madison, Wisconsin, USA; Telethon Kids Institute, University of Western Australia, Perth, Western Australia, Australia; Airway Disease Infection Section and MRC & Asthma UK Centre in Allergic Mechanisms of Asthma, National Heart and Lung Institute, Imperial College London, Norfolk Place, London, United Kingdom; Division of Infection, Immunity and Respiratory Medicine, The University of Manchester; Child Health Research Centre, The University of Queensland, Brisbane, Queensland, Australia; Department of Biochemistry and Molecular Biology, The University of Melbourne, Parkville, Victoria, Australia; Department of Pathology, The University of Melbourne, Parkville, Victoria, Australia

## Abstract

Events in early life contribute to subsequent risk of asthma; however, the causes and trajectories of childhood wheeze are heterogeneous and do not always result in asthma. Similarly, not all atopic individuals develop wheeze, and vice versa. The reasons for these differences are unclear. Using unsupervised model-based cluster analysis, we identified latent clusters within a prospective birth cohort with deep immunological and respiratory phenotyping. We characterised each cluster in terms of immunological profile and disease risk, and replicated our results in external cohorts from the UK and USA. We discovered three distinct trajectories, one of which is a high-risk “atopic” cluster with increased propensity for allergic diseases throughout childhood. Atopy contributes varyingly to later wheeze depending on cluster membership. Our findings demonstrate the utility of unsupervised analysis in elucidating heterogeneity in asthma pathogenesis and provide a foundation for improving management and prevention of childhood asthma.

## 1 Introduction

Asthma is a global health problem, and there is a pressing need for better understanding of its pathogenesis [1]. Both genetic and environmental factors are involved in asthma [2, 3], and the “hygiene hypothesis” proposes that modern changes to hygiene, sanitation and living environment have modified human exposures to microbes, with subsequent effects on early-life immune development [4]. However, the clinical presentation and prognosis of childhood wheeze is highly variable: some children remit; others remit but relapse in later life; and yet others have wheeze persisting into adult asthma [5]. These differences suggest that the underlying causes of disease also differ from person to person. For example, while asthma is commonly linked to allergy, not all individuals with wheeze are sensitised to allergen, and vice versa [6]. As such, childhood asthma is a heterogeneous condition [7, 8], and this greatly complicates the study of its pathogenesis [9]. We postulate that there are subpopulations in early childhood, each sharing similar patterns of pathophysiology, disease susceptibility and phenotype that permit categorisation into clusters. If we can agnostically identify these clusters, then we may identify the biological mechanisms that underlie them, and find targets for early intervention that are specific for different asthma subtypes.

Older attempts at subtyping asthma susceptibility relied on supervised classification, using expert knowledge and cut-offs to define clusters. For example, specific immunoglobulin E (IgE) ≥ 0.35 kU/L; wheal diameter ≥ 3 mm in a skin prick test (SPT); or symptom score surpassing a threshold – would determine classification into a high-risk profile [10, 11]. However, these cut-offs vary with age, gender or other parameters, and may not accurately reflect true attribution of risk [12]. Hence, they often continue to produce heterogeneous groups. Furthermore, previous studies tended to focus on a single “domain”, for instance grouping only by immunological response [13], symptomatology or timing of disease [14, 15]. Recently, researchers have turned to unsupervised approaches, such as model-based cluster analysis and latent class analysis (LCA) [16–21]. These do not require experts to supply cut-offs, but can instead “learn” boundaries from the data. They can potentially uncover patterns of similarity not immediately obvious to the human eye. Finally, these methods can cover a broader range of domains, incorporating measurements from multiple sources to determine clusters that are potentially informative of asthma risk.

Here, we use a data-driven unsupervised framework together with a comprehensively-phenotyped birth cohort (the Childhood Asthma Study, CAS) to define developmental trajectories during preschool years, a period known to be critical to asthma pathogenesis. Specifically, we 1) discover, using non-parametric mixture models, latent clusters that define early childhood trajectories of immune function and susceptibility to respiratory infection; 2) investigate how these clusters relate to differential profiles of asthma susceptibility, and to existing definitions of atopy; 3) identify risk factors for asthma within each cluster, which may differ across said clusters; 4) summarise and simplify our findings in decision trees; and 5) externally validate the clusters in independent cohorts.

## 2 Results

Our discovery dataset was CAS, a Western Australian birth cohort (*N*=263) enriched for parental asthma history [22], with clinical, immunological, and respiratory measurements from the first ten years of life (**Methods**). Clinical variables included demographics, incidence of allergic disease and family history. Immunological variables included IgE, IgG4, and IgG antibody levels for common allergens, as well as a combination antibody assay (Phadiatop) which covers multiple allergens [23]. We measured frequency and severity of respiratory infections, and performed 16S rRNA sequencing on nasopharyngeal aspirates (NPAs) collected during respiratory infections (disease samples) or routine check-ups (healthy samples). These NPAs have been classified by Teo et al [24, 25], based on clustering of microbial composition, into microbiome profile groups (MPGs) that were associated with healthy respiratory states (health-associated MPGs, e.g. *Alloiococcus-, Staphylococcus*- or *Corynebacterium*-dominated) or infectious respiratory states (infection-associated MPGs, e.g. *Moraxella-, Haemophilus-*, or *Streptococcus*-dominated).

To identify latent clusters, we applied non-parametric expectation-maximisation (EM) mixture modelling (“npEM”) to CAS. We used a largely non-selective approach to the choice of features, but we did explicitly exclude variables with excessive missing data (**Methods, Supplementary Methods**), as well as primary outcomes such as yearly incidence of wheeze and physician-diagnosed asthma (**Supplementary Table 1**). By virtue of study design and exclusion criteria, most included variables were related to immunological function or respiratory infection in the first three years of life. Individuals were assigned to a cluster if the mixture model determined ≥ 90% probability of membership in that cluster (**Supplementary Methods**). We described each cluster in terms of key characteristics and significant cluster-specific predictors for age-five wheeze. We also built an npEM-derived classifier to cluster samples with low missing data, and classify individuals from two comparable datasets for replication - the Manchester Asthma and Allergy Study (MAAS) (#=1085) [26] from Manchester, UK, and the Childhood Origins of Asthma Study (COAST) (*N*=289) from Wisconsin, USA [27] (**Supplementary Table 2; Supplementary Figure 1**). Finally, we developed a decision tree classifier, which allowed us to simplify cluster description and determine which features best separated clusters.

Using npEM-based clustering and classification in CAS, we identified three distinct clusters from 217 individuals and 174 clustering features (**Figure 1**): low-risk CAS1 (*N*=88, 25% wheeze at age 5), low-risk but allergy-susceptible CAS2 (*N*=107, 21% wheeze at age 5) and high-risk CAS3 (*N*=22, 76% wheeze at age 5) (**Figure 2**). Forty-six individuals in CAS had excessive missing data and were not classifiable (**Methods**).

**Figure 1:**
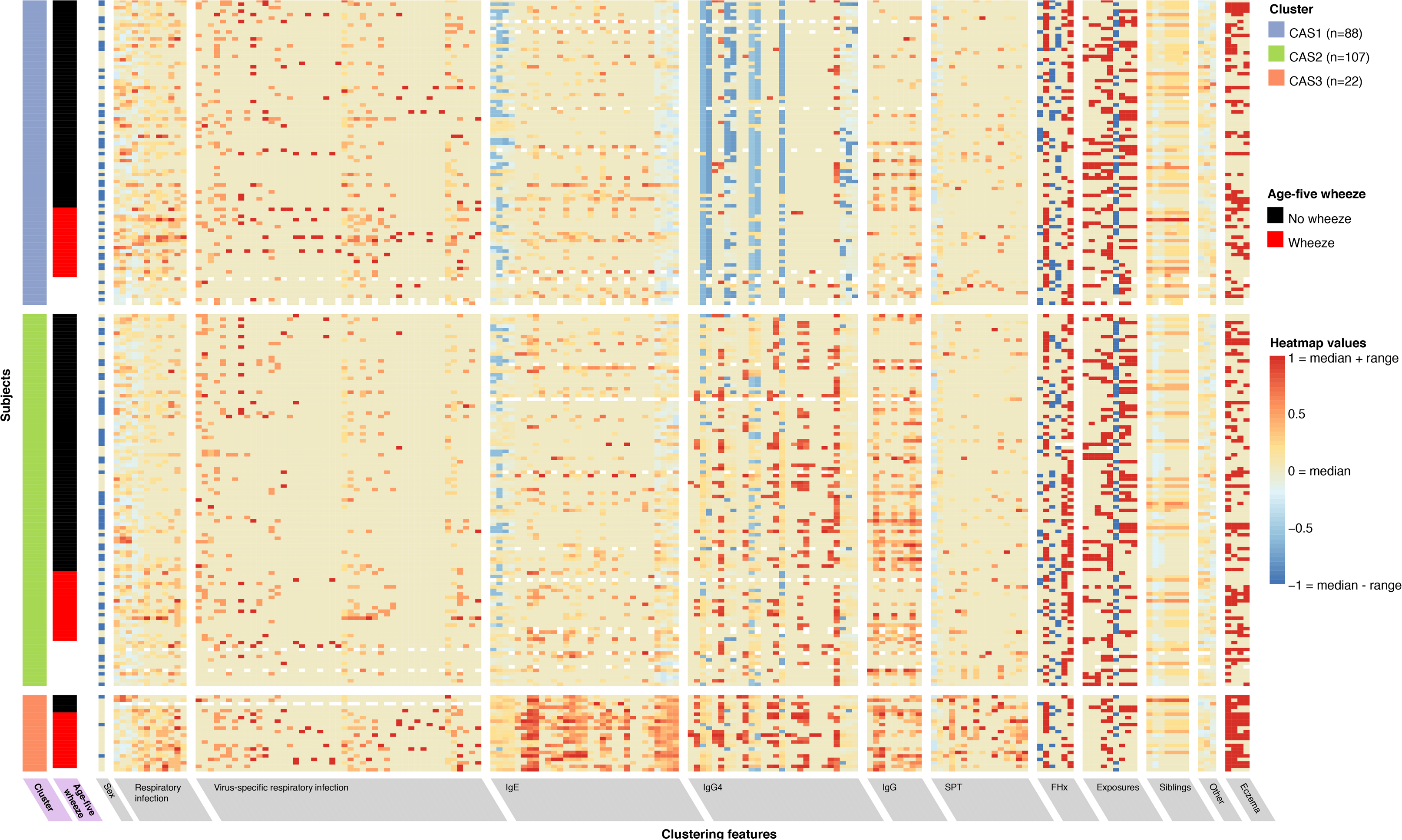
Non-parametric mixture-model based clustering of CAS dataset, based on 174 features. SPT = skin prick test. White spaces within the heatmap indicate missing data. Rows represent individuals; columns represent clustering features with general categories as labelled on grey background. Variables with grey background are clustering features ordered by category or type of variable first (e.g. all HDM IgE-related variables grouped together), then by timepoint (earlier to later, from left to right). Variables with lilac background indicate resultant cluster membership and outcome variable (age-five wheeze). Heatmap values are scaled relative to range and median values for each feature; the median is coloured beige-yellow, the median + range red, and median < range blue. For sex, -1/blue = female, 0/yellow (median) = male.

**Figure 2:**
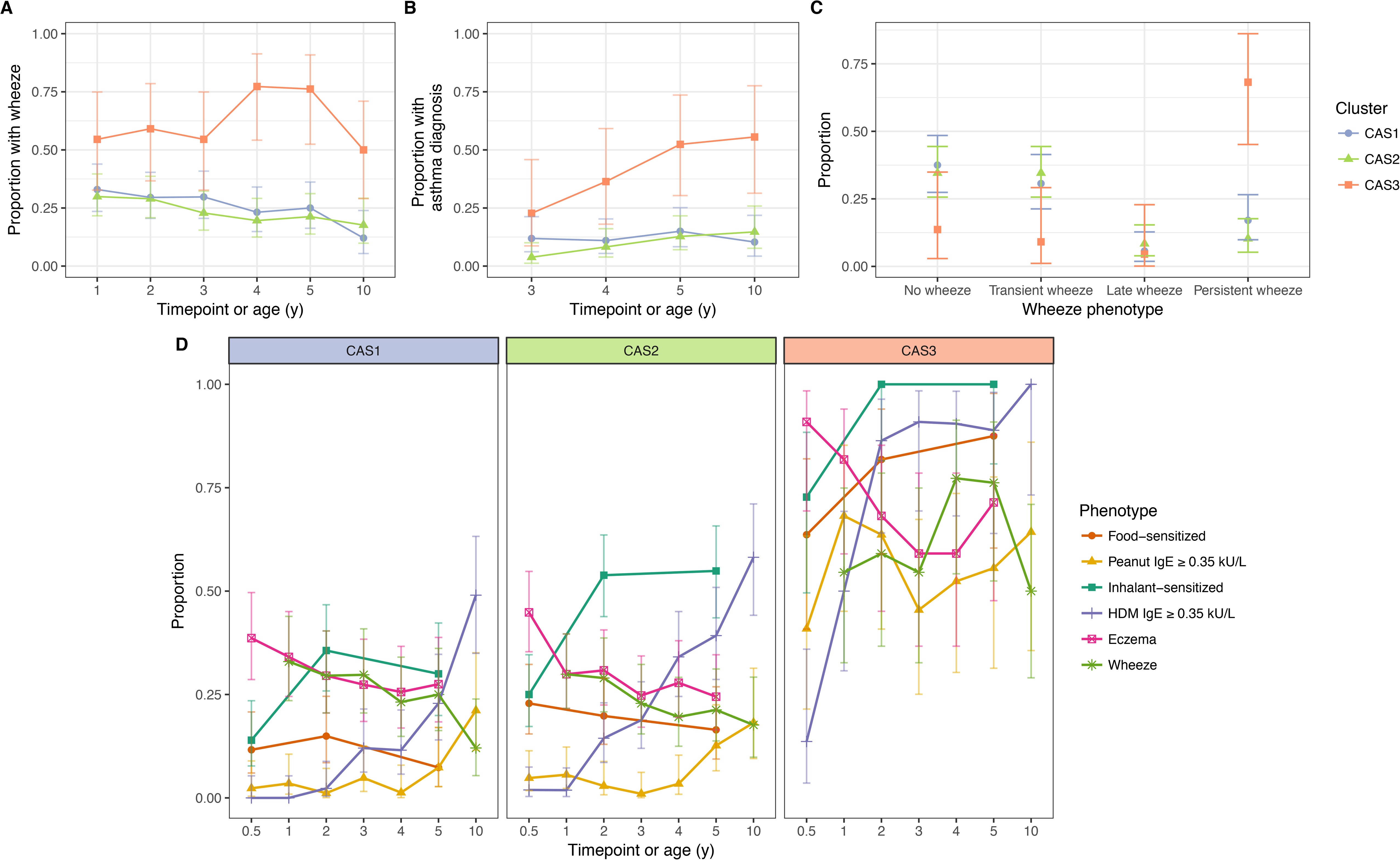
Incidence of multiple phenotypes, including parent-reported wheeze (A), physician-diagnosed asthma (B), defined wheeze phenotypes (C), in relation to food and inhalant sensitisation (D), stratified by cluster and time in the CAS dataset. Points indicate observed proportion; bars indicate 95% CI (binomial distribution). Wheeze phenotypes defined as: no wheeze = no wheeze at ages 1 to 3, or age 5; transient wheeze = any wheeze at ages 1 to 3, but not age 5; late wheeze = wheeze at age 5, but not ages 1 to 3; persistent wheeze = any wheeze at both ages 1 to 3 and age 5. Food sensitization defined as peanut IgE ≥ 0.35 kU/L at any age, or cow’s milk, egg white, peanut SPT > 2 or 3 mm for age ≤ 2 or > 2 respectively. Inhalant sensitization defined as HDM, cat, couchgrass, ryegrass, mould or Phadiatop IgE ≥ 0.35 kU/L at any age, or mould SPT *(Alternaria* or *Aspergillus* spp.) > 2 or 3 mm for age ≤ 2 or > 2 respectively.

### 2.1 CAS1: low-risk, non-atopic cluster with transient wheeze

CAS1 was a low-risk cluster with infrequent and transient respiratory wheeze. Rates of wheeze declined from 33% at age 1 to 12% by age 10 (**Table 1; Figure 2**). In this cluster, Th2 cytokine responses of PBMCs to allergen stimulation were minimal; and rates of allergen sensitisation (as measured by IgE or SPT) were the lowest among all groups (**Table 2, Supplementary Tables 3B-D**; Figure 3). IgG and IgG4 were also low across all allergens.

**Table 1:** Comparison of selected demographic and clinical variables in CAS clusters

| Variable | Age (y) | CAS1 (N=88) | CAS2 (N=107) | CAS3 (N=22) | P-value (unadjusted) |  |  |  | Feature? |
| --- | --- | --- | --- | --- | --- | --- | --- | --- | --- |
|  |  | Prop. (95% CI) | Prop. (95% CI) | Prop. (95% CI) | Overall | 1 vs. 2 | 1 vs. 3 | 2 vs. 3 |  |
| Sex = male |  | 55% (44%-65%) | 51% (42%-61%) | 86% (71%-100%) | **7.3E-03** | 0.67 | **6.8E-03** | **3.7E-03** | **Yes** |
| Maternal asthma |  | 51% (40%-62%) | 41% (32%-51%) | 59% (37%-81%) | 0.19 | 0.19 | 0.63 | 0.16 | **Yes** |
| Paternal asthma |  | 22% (13%-30%) | 44% (35%-54%) | 23% (3.7%-42%) | **2.2E-03** | **1.3E-03** | 1 | *0.093* | **Yes** |
| Wheeze | 1 | 33% (23%-43%) | 30% (21%-39%) | 55% (32%-77%) | *0.092* | 0.76 | *0.084* | **0.046** | No |
|  | 5 | 25% (15%-35%) | 21% (13%-30%) | 76% (56%-96%) | **7.1E-06** | 0.59 | **2.6E-05** | **3.4E-06** | No |
|  | 10 | 12% (3.4%-21%) | 18% (8.4%-27%) | 50% (24%-76%) | **3.1E-03** | 0.46 | **1.5E-03** | **0.011** | No |
| Asthma | 5 | 15% (7%-23%) | 13% (5.9%-20%) | 52% (29%-76%) | **4.1E-04** | 0.83 | **7.7E-04** | **2.1E-04** | No |
|  | 10 | 10% (2.3%-18%) | 15% (6.1%-23%) | 56% (30%-81%) | **2.6E-04** | 0.59 | **1.8E-04** | **7.9E-04** | No |
| Eczema | 6m | 39% (28%-49%) | 45% (35%-54%) | 91% (78%-100%) | **2.4E-05** | 0.47 | **7.9E-06** | **9.0E-05** | **Yes** |
|  | 1 | 34% (24%-44%) | 30% (21%-39%) | 82% (64%-99%) | **2.5E-05** | 0.54 | **7.2E-05** | **1.4E-05** | **Yes** |
|  | 5 | 28% (18%-37%) | 24% (16%-33%) | 71% (50%-92%) | **2.1E-04** | 0.73 | **3.3E-04** | **7.9E-05** | No |
| Atopic rhinoconjunctivitis | 5 | 30% (20%-40%) | 39% (29%-49%) | 76% (56%-96%) | **6.4E-04** | 0.21 | **2.7E-04** | **3.2E-03** | No |
|  |  | Mean (95% CI) | Mean (95% CI) | Mean (95% CI) | Overall | 1 vs. 2 | 1 vs. 3 | 2 vs. 3 |  |
| BMI (kg/m<sup>2</sup>) | 3 | 16 (16-17) | 16 (16-17) | 16 (16-17) | 0.86 | 0.65 | 0.68 | 0.8 | No* |
|  | 4 | 16 (16-17) | 16 (16-16) | 17 (16-17) | 0.59 | 0.76 | 0.32 | 0.39 | No |
|  | 5 | 16 (16-16) | 16 (16-16) | 16 (15-17) | 0.71 | 0.56 | 0.48 | 0.67 | No |
|  | 10 | 18 (17-19) | 18 (17-18) | 18 (17-19) | 0.89 | 0.75 | 1 | 0.62 | No |
| Number of older siblings | 0 | 0.93 (0.72-1.1) | 0.53 (0.38-0.69) | 0.77 (0.32-1.2) | **4.5E-03** | **1.0E-03** | 0.37 | 0.25 | **Yes** |
|  | 2 | 0.85 (0.66-1) | 0.5 (0.34-0.65) | 0.77 (0.32-1.2) | **2.8E-03** | **6.5E-04** | 0.48 | 0.16 | **Yes** |
|  | 5 | 0.68 (0.5-0.85) | 0.39 (0.25-0.54) | 0.67 (0.23-1.1) | **0.016** | **5.1E-03** | 0.75 | 0.12 | No |
|  |  | Geom. mean (95% CI) | Geom. mean (95% CI) | Geom. mean (95% CI) | Overall | 1 vs. 2 | 1 vs. 3 | 2 vs. 3 |  |
| Vitamin D (nmol/L) | 1 | 60 (55-64) | 59 (55-63) | 59 (52-67) | 0.93 | 0.98 | 0.76 | 0.7 | No |
|  | 2 | 57 (54-61) | 58 (55-61) | 47 (40-55) | **0.012** | 0.82 | **5.4E-03** | **4.4E-03** | No |
|  | 5 | 89 (83-95) | 84 (79-89) | 77 (69-84) | *0.057* | 0.46 | **0.016** | *0.056* | No |

BMI = body mass index; feature? = whether variable was used as a clustering feature or not; geom. mean = geometric mean; prop. = proportion. For categorical variables, associations were tested using Fisher exact test; for continuous variables, Kruskal-Wallis and Mann-Whitney-Wilcoxon. Bold text indicates statistical significance ( $p < 0.05$ ); italics indicate near-significance ( $p < 0.10$ ). \*Not used as clustering feature, as BMI is a derived variable. Height and weight at age 3 were used instead.

**Table 2:**
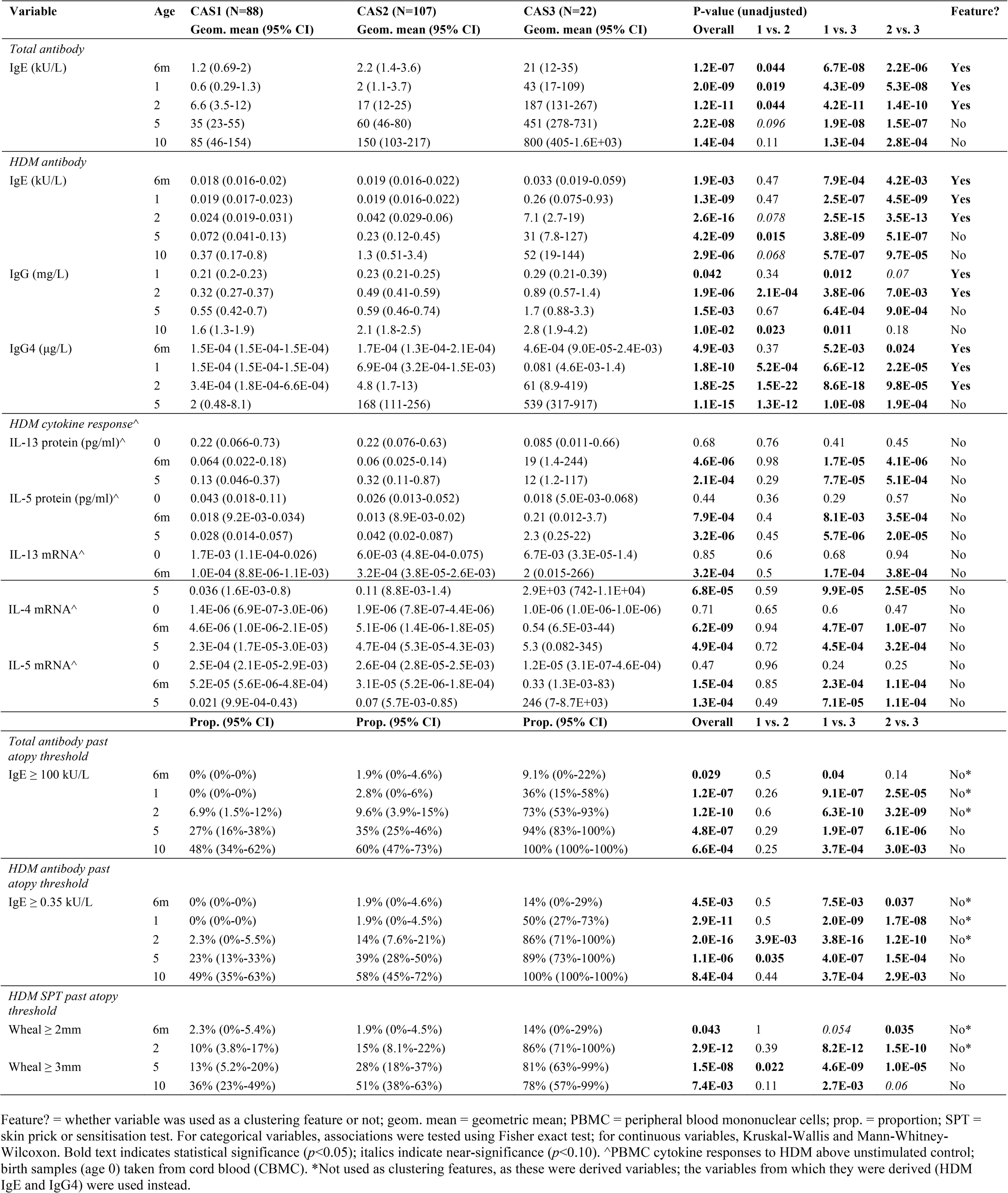
Comparison of HDM-associated immunological variables in CAS clusters

Frequency of respiratory infection in CAS1 were intermediate to low (**Table 3**). However, in this cluster, a high frequency of lower respiratory infections (LRIs) in childhood, especially wheezy LRIs (wLRIs), was a risk factor for age-five wheeze - even after adjusting for sex, BMI and parental history of asthma as demographic covariates (**Table 4; Figure 4A-B**). After multiple regression analysis with stepwise backward elimination (**Methods**), three variables remained significant in the one model: age-three wLRI frequency (odds ratio OR 8.3 per unit increase, *p*=3.4×10^−2^); age-four LRI frequency (OR 3.6, *p*=0.022); and proportion of infection-associated MPGs (*Streptococcus*, *Haemophilus*, *Moraxella*) in age-two-to-four healthy NPAs (OR 0.13 per quartile, *p*=0.016). Repeated-measures ANOVA confirmed that LRI and wLRI frequency in the first 3 years of life were predictors for age-five wheeze within CAS1 (**Supplementary Table 4**).

**Table 3:** Comparison of selected respiratory disease-related variables in CAS clusters

| Variable | Age (y) | CAS1 (N=88) | CAS2 (N=107) | CAS3 (N=22) | P-value (unadjusted) |  |  |  | Feature? |
| --- | --- | --- | --- | --- | --- | --- | --- | --- | --- |
|  |  | Mean (95% CI) | Mean (95% CI) | Mean (95% CI) | Overall | 1 vs. 2 | 1 vs. 3 | 2 vs. 3 |  |
| Any ARI (events per y) | 1 | 4.4 (3.9-4.9) | 3.6 (3.1-4.1) | 4.5 (3.3-5.6) | **0.044** | **0.018** | 0.86 | 0.16 | No* |
|  | 2 | 4.5 (3.8-5.2) | 3.6 (3.2-4) | 4.7 (3.4-6) | 0.13 | 0.13 | 0.56 | **0.077** | No* |
|  | 3 | 3.7 (3.1-4.3) | 3.4 (3-3.9) | 4 (2.7-5.4) | 0.74 | 0.56 | 0.77 | 0.53 | No* |
|  | 4 | 3 (2.4-3.6) | 2.7 (2.2-3.2) | 3.7 (2.4-4.9) | 0.28 | 0.48 | 0.28 | 0.11 | No |
|  | 5 | 2 (1.5-2.5) | 1.9 (1.5-2.3) | 1.5 (0.9-2.1) | 0.94 | 0.83 | 0.89 | 0.74 | No |
| URI (events per y) | 1 | 2.9 (2.4-3.3) | 2.6 (2.2-3) | 2.5 (1.7-3.3) | 0.59 | 0.34 | 0.5 | 0.96 | **Yes** |
|  | 2 | 3.2 (2.6-3.7) | 2.6 (2.2-3) | 2.5 (1.2-3.8) | 0.19 | 0.19 | 0.12 | 0.34 | **Yes** |
|  | 3 | 2.7 (2.2-3.2) | 2.8 (2.4-3.3) | 2.2 (1.3-3.2) | 0.45 | 0.41 | 0.59 | 0.24 | **Yes** |
|  | 4 | 2.1 (1.7-2.6) | 2.2 (1.8-2.7) | 1.7 (0.77-2.7) | 0.5 | 0.94 | 0.26 | 0.27 | No |
|  | 5 | 1.6 (1.1-2) | 1.5 (1.2-1.9) | 0.67 (0.2-1.1) | *0.081* | 0.76 | **0.047** | **0.026** | No |
| LRI (events per y) | 1 | 1.6 (1.2-1.9) | 0.98 (0.76-1.2) | 2 (1.3-2.6) | **4.0E-03** | **0.021** | **0.17** | **2.6E-03** | **Yes** |
|  | 2 | 1.4 (0.98-1.7) | 1 (0.81-1.2) | 2.2 (1.6-2.9) | **2.5E-03** | 0.83 | **6.1E-03** | **2.0E-04** | **Yes** |
|  | 3 | 1 (0.76-1.3) | 0.6 (0.4-0.8) | 1.8 (1.1-2.6) | **6.1E-04** | **0.02** | **0.039** | **2.7E-04** | **Yes** |
|  | 4 | 0.87 (0.52-1.2) | 0.46 (0.3-0.63) | 2 (1.1-2.8) | **1.7E-05** | 0.3 | **3.5E-04** | **1.6E-06** | No |
|  | 5 | 0.42 (0.24-0.6) | 0.36 (0.24-0.48) | 0.86 (0.44-1.3) | **0.019** | 1 | **0.011** | **7.5E-03** | No |
| Wheezy LRI (wLRI, events per y) | 1 | 0.47 (0.3-0.63) | 0.24 (0.15-0.34) | 0.64 (0.19-1.1) | *0.054* | **0.036** | 0.61 | *0.065* | **Yes** |
|  | 2 | 0.68 (0.45-0.91) | 0.41 (0.26-0.56) | 1 (0.56-1.5) | **5.2E-03** | *0.063* | *0.066* | **1.7E-03** | **Yes** |
|  | 3 | 0.59 (0.37-0.81) | 0.3 (0.17-0.44) | 1.4 (0.78-2.1) | **4.6E-05** | *0.065* | **2.5E-03** | **6.6E-06** | **Yes** |
|  | 4 | 0.52 (0.25-0.79) | 0.32 (0.18-0.46) | 1.9 (0.95-2.8) | **4.5E-08** | 0.86 | **9.3E-07** | **3.3E-08** | No |
|  | 5 | 0.28 (0.13-0.42) | 0.23 (0.13-0.33) | 0.76 (0.36-1.2) | **2.3E-03** | 0.99 | **2.0E-03** | **1.2E-03** | No |
| Febrile LRI (fLRI, events per y) | 1 | 0.36 (0.22-0.51) | 0.28 (0.16-0.4) | 0.55 (0.28-0.81) | **0.025** | 0.24 | *0.071* | **6.4E-03** | **Yes** |
|  | 2 | 0.36 (0.23-0.5) | 0.33 (0.22-0.43) | 0.95 (0.46-1.4) | **0.01** | 1 | **6.1E-03** | **3.8E-03** | **Yes** |
|  | 3 | 0.38 (0.21-0.55) | 0.16 (0.09-0.23) | 0.52 (0.13-0.92) | *0.06* | *0.063* | 0.44 | **0.04** | **Yes** |
|  | 4 | 0.3 (0.13-0.47) | 0.15 (0.064-0.24) | 0.43 (0.16-0.7) | **0.021** | 0.18 | *0.091* | **4.9E-03** | No |
|  | 5 | 0.19 (0.082-0.3) | 0.14 (0.06-0.21) | 0.19 (0-0.42) | 0.83 | 0.55 | 0.91 | 0.8 | No |
| Severe LRI (wLRI or fLRI, events per y) | 1 | 0.69 (0.5-0.89) | 0.44 (0.29-0.58) | 1 (0.49-1.5) | **0.012** | **0.027** | 0.25 | **9.1E-03** | No* |
|  | 2 | 0.9 (0.62-1.2) | 0.59 (0.43-0.75) | 1.6 (1.1-2.2) | **7.9E-04** | 0.22 | **5.2E-03** | **1.2E-04** | No* |
|  | 3 | 0.73 (0.49-0.97) | 0.37 (0.23-0.51) | 1.5 (0.85-2.2) | **1.6E-04** | **0.032** | **0.01** | **3.8E-05** | No* |
|  | 4 | 0.63 (0.32-0.94) | 0.36 (0.21-0.52) | 1.9 (1-2.8) | **2.8E-07** | 0.56 | **5.9E-06** | **8.4E-08** | No |
|  | 5 | 0.36 (0.19-0.53) | 0.27 (0.17-0.38) | 0.76 (0.36-1.2) | **0.015** | 0.88 | **0.012** | **5.0E-03** | No |

|  |  | Prop. (95% CI) | Prop. (95% CI) | Prop. (95% CI) | Overall | 1 vs. 2 | 1 vs. 3 | 2 vs. 3 |  |
| --- | --- | --- | --- | --- | --- | --- | --- | --- | --- |
| >20% *Streptococcus* in first infection-naïve NPA sample | 7w | 11% (0.34%-23%) | 15% (3.3%-26%) | 44% (3.9%-85%) | *0.081* | 0.75 | **0.042** | *0.065* | No |
|  | 6m | 7.6% (1.6%-14%) | 18% (10%-26%) | 14% (0%-31%) | 0.12 | **0.045** | 0.39 | 1 | No |
| % Healthy NPAs with infection-associated MPGs | 0-2 | 49% (38%-59%) | 32% (24%-39%) | 62% (47%-76%) | **1.2E-03** | **0.013** | 0.2 | **5.5E-04** | No |
|  | 2-4 | 46% (37%-55%) | 44% (37%-51%) | 45% (29%-61%) | 0.9 | 0.67 | 0.92 | 0.8 | No |

**Table 4:** Analysis of selected predictors for age-five wheeze within each CAS cluster, with demographic covariates (sex, BMI, parental history of asthma)

| Selected predictors for age-five wheeze |  | CAS1 (N=88) |  | CAS2 (N=107) |  | CAS3 (N=22) |  | All (N=261) |  |
| --- | --- | --- | --- | --- | --- | --- | --- | --- | --- |
|  |  | OR (95% CI) | P-value | OR (95% CI) | P-value | OR (95% CI) | P-value | OR (95% CI) | P-value |
| ARI (events per y) | 1 | 1.1 (0.88-1.5) | 0.36 | 1.1 (0.87-1.3) | 0.51 | 0.57 (0.29-0.93) | **0.046** | 1 (0.89-1.2) | 0.76 |
|  | 2 | 1.1 (0.94-1.3) | 0.22 | 1 (0.81-1.3) | 0.82 | 0.43 (0.077-0.89) | 0.12 | 1 (0.93-1.2) | 0.44 |
|  | 3 | 1.1 (0.87-1.3) | 0.58 | 1.1 (0.91-1.4) | 0.3 | 0.67 (0.36-1) | 0.1 | 1 (0.93-1.2) | 0.48 |
|  | 4 | 1.2 (0.99-1.4) | *0.074* | 1.2 (1-1.5) | **0.032** | 0.63 (0.27-1.1) | 0.15 | 1.2 (1-1.3) | **0.013** |
| LRI (events per y) | 1 | 0.97 (0.71-1.3) | 0.84 | 1 (0.61-1.5) | 0.99 | 0.48 (0.13-1.1) | 0.16 | 1 (0.81-1.2) | 0.92 |
|  | 2 | 1.2 (0.88-1.6) | 0.26 | 1.5 (0.97-2.5) | *0.069* | 0.99 (0.34-2.6) | 0.98 | 1.4 (1.1-1.7) | **5.3E-03** |
|  | 3 | 2 (1.3-3.2) | **2.3E-03** | 2.6 (1.5-5.3) | **2.7E-03** | 0.98 (0.4-2.6) | 0.96 | 2 (1.5-2.7) | **3.8E-06** |
|  | 4 | 2 (1.4-3.4) | **2.0E-03** | 3.6 (1.8-8.3) | **6.5E-04** | 1.9 (0.57-8.4) | 0.32 | 2.5 (1.8-3.6) | **1.5E-07** |
| Wheezy LRI (events per y) | 1 | 1.3 (0.68-2.4) | 0.43 | 1.1 (0.35-3) | 0.83 | 2.6 (0.62-58) | 0.34 | 1.5 (0.98-2.3) | *0.06* |
|  | 2 | 1.2 (0.8-2) | 0.33 | 1.6 (0.89-2.9) | 0.12 | 2.4 (0.67-16) | 0.24 | 1.6 (1.2-2.2) | **5.6E-03** |
|  | 3 | 2.8 (1.6-5.6) | **1.3E-03** | 3 (1.4-8) | **0.016** | 1.2 (0.43-4.6) | 0.76 | 2.7 (1.8-4.2) | **4.1E-06** |
|  | 4 | 2.5 (1.5-5) | **4.0E-03** | 6.3 (2.5-21) | **6.8E-04** | 7.1 (1.2-169) | 0.1 | 3.9 (2.5-6.7) | **5.4E-08** |
| Febrile LRI (events per y) | 1 | 1.6 (0.77-3.6) | 0.21 | 0.84 (0.28-1.9) | 0.71 | 7.3 (0.78-178) | 0.12 | 1.5 (0.93-2.4) | *0.098* |
|  | 2 | 1 (0.44-2.2) | 1 | 4.8 (1.8-15) | **3.9E-03** | 1.6 (0.48-10) | 0.5 | 2.3 (1.4-3.9) | **1.2E-03** |
|  | 3 | 2 (1-4.8) | *0.08* | 4.3 (1.2-15) | **0.02** | 4.2 (0.55-519) | 0.37 | 2.4 (1.4-4.3) | **2.3E-03** |
|  | 4 | 1.8 (0.97-4.1) | *0.092* | 2.6 (0.88-8.3) | *0.082* | 1.1 (0.11-18) | 0.93 | 2.2 (1.3-4) | **5.9E-03** |
| % Healthy NPAs with infection-associated MPGs | 0-2 | 0.9 (0.13-5.7) | 0.91 | 2.6 (0.43-16) | 0.3 | NA | NA | 2.3 (0.79-6.7) | 0.13 |
|  | 2-4 | 0.086 (6.8E-03-0.71) | **0.034** | 0.8 (0.077-7.5) | 0.85 | 4.4E+03 (2.1-2.5E+12) | 0.13 | 0.49 (0.14-1.6) | 0.24 |
| Quartile of % healthy NPAs with infection-associated MPGs | 0-2 | 1 (0.54-1.8) | 0.98 | 1.3 (0.72-2.4) | 0.36 | NA | NA | 1.3 (0.89-1.8) | 0.19 |
|  | 2-4 | 0.45 (0.19-0.88) | **0.035** | 1 (0.51-2.1) | 0.9 | NA | NA | 0.8 (0.53-1.2) | 0.24 |
| HDM IgE (kU/L)* | 6m | 8 (0.85-94) | *0.074* | 0.93 (0.14-3.6) | 0.92 | 3.4 (0.26-180) | 0.4 | 2.3 (0.99-5.8) | *0.054* |
|  | 1 | 1.5 (0.22-7.8) | 0.65 | 0.54 (0.039-2.3) | 0.51 | 39 (2.5-22000) | *0.082* | 2.7 (1.5-5) | **0.00089** |
|  | 2 | 0.93 (0.28-2.5) | 0.89 | 2 (1.2-3.7) | **0.016** | 1.4 (0.38-4.8) | 0.62 | 2 (1.5-2.8) | **2.80E-05** |
|  | 3 | 1.4 (0.68-2.9) | 0.32 | 1.5 (0.9-2.4) | 0.12 | 1.5 (0.4-5.2) | 0.55 | 1.7 (1.3-2.2) | **1.00E-04** |
|  | 4 | 1.9 (0.94-4.1) | *0.086* | 1.9 (1.2-3.1) | **0.011** | 1.4 (0.31-5.5) | 0.64 | 1.9 (1.5-2.5) | **3.70E-06** |
| Peanut IgE (kU/L)* | 6m | 2.5 (0.78-9) | 0.13 | 1.5 (0.54-3.8) | 0.41 | 1.1 (0.3-3.7) | 0.92 | 2.3 (1.4-3.9) | **0.0014** |
|  | 1 | 1.7 (0.48-6.3) | 0.39 | 2.2 (0.65-6.9) | 0.19 | 0.47 (0.095-1.6) | 0.27 | 2.2 (1.4-3.6) | **0.00098** |
|  | 2 | 0.51 (0.097-2) | 0.37 | 3 (0.74-12) | 0.12 | 2 (0.51-13) | 0.37 | 2.7 (1.6-4.9) | **0.00046** |
|  | 3 | 1.7 (0.46-5.5) | 0.37 | 0.53 (0.015-3.8) | 0.61 | 3.3 (0.94-26) | 0.13 | 2.6 (1.6-4.8) | **0.00068** |
|  | 4 | 0.2 (0.00073-2.9) | 0.36 | 0.96 (0.19-3.2) | 0.95 | 1.4 (0.49-6.5) | 0.54 | 2.1 (1.3-3.7) | **0.006** |
| Cat IgE (kU/L)* | 6m | 6.6 (0.77-61) | 0.079 | 2.2 (0.62-7.6) | 0.2 | 0.24 (0.012-3.2) | 0.29 | 2.3 (0.96-5.4) | 0.061 |
|  | 1 | 2.1 (0.13-30) | 0.57 | 4 (0.54-32) | 0.16 | 0.45 (0.053-2.8) | 0.41 | 3.5 (1.4-9.5) | **0.0099** |
|  | 2 | 0.55 (0.042-3.7) | 0.57 | 2.1 (0.59-7) | 0.22 | 2.2 (0.42-26) | 0.42 | 2.6 (1.3-5.5) | **0.0065** |
|  | 3 | 1.7 (0.49-5.6) | 0.35 | 1.4 (0.21-6.7) | 0.66 | 1.3 (0.29-6.9) | 0.77 | 2.5 (1.3-4.9) | **0.0065** |
|  | 4 | 0.75 (0.0088-13) | 0.86 | 1.5 (0.53-3.9) | 0.4 | 0.83 (0.17-4.4) | 0.81 | 2.4 (1.3-4.8) | **0.006** |
| Couch grass IgE (kU/L)* | 6m | 2.8 (0.51-14) | 0.21 | 1.3 (0.3-4.5) | 0.68 | 0.98 (0.048-59) | 0.99 | 1.7 (0.71-3.9) | 0.22 |
|  | 1 | 0.38 (0.017-2.8) | 0.42 | 0.33 (0.01-2.9) | 0.41 | 0.15 (0.0058-1.5) | 0.14 | 0.63 (0.19-1.7) | 0.4 |
|  | 2 | 0.085 (0.0034-0.7) | 0.057 | 1.1 (0.14-6.3) | 0.9 | 25 (1.6-1100) | **0.046** | 2.1 (0.99-4.7) | 0.053 |
|  | 3 | 2 (0.44-8) | 0.29 | 6.1e-06 (NA-8.1e+54) | 0.99 | 2.3 (0.57-14) | 0.29 | 2.5 (1.3-5.1) | **8.90E-03** |
|  | 4 | 8.4e-13 (NA-3.5e+172) | 0.99 | 1.6 (0.55-4.1) | 0.34 | 1.9 (0.54-10) | 0.35 | 2 (1.3-3.4) | **4.30E-03** |
| Phadiatop IgE (PAU/L)* | 6m | 1.2 (0.44-2.9) | 0.73 | 1.3 (0.65-2.6) | 0.43 | 2.2 (0.66-12) | 0.25 | 2 (1.3-2.9) | **0.00078** |
|  | 1 | 0.73 (0.2-2.5) | 0.63 | 1.1 (0.41-2.8) | 0.85 | 1.6 (0.23-18) | 0.67 | 2.1 (1.3-3.4) | **0.0021** |
|  | 2 | 0.33 (0.091-1) | 0.065 | 2.1 (0.81-5.9) | 0.13 | 2.5 (0.18-70) | 0.52 | 2 (1.3-3) | **0.0012** |
|  | 3 | 1.8 (0.8-4) | 0.16 | 1.4 (0.72-2.8) | 0.31 | 8.4 (0.53-380) | 0.19 | 2 (1.4-2.9) | **8.00E-05** |
|  | 4 | 1.8 (0.91-3.8) | 0.094 | 2.4 (1.3-4.8) | **0.01** | 2.7 (0.16-66) | 0.5 | 2.2 (1.6-3.2) | **2.20E-06** |
| HDM IgG4 (µg/L)* | 6m | NA (NA-NA) | 0.55 | 0.053 (NA-6.5e+24) | 0.99 | 28 (1.7e-34-NA) | 0.99 | 1.4 (0.88-2.6) | 0.17 |
|  | 1 | NA (NA-NA) | 0.61 | 1.1 (0.8-1.5) | 0.5 | 0.9 (0.58-1.3) | 0.6 | 1.2 (1-1.4) | 0.053 |
|  | 2 | 1.1 (0.71-1.6) | 0.67 | 1.1 (0.85-1.4) | 0.61 | 0.4 (0.038-1.2) | 0.26 | 1.1 (1-1.3) | 0.056 |
|  | 3 | 1.1 (0.85-1.5) | 0.35 | 1.1 (0.77-2) | 0.64 | 0.94 (0.19-2.3) | 0.9 | 1.1 (0.98-1.2) | 0.1 |
|  | 4 | 1.2 (0.98-1.5) | 0.082 | 0.89 (0.7-1.1) | 0.33 | 0.46 (0.031-5.4) | 0.53 | 1.1 (1-1.3) | **0.034** |
| Peanut IgG4 (µg/L)* | 6m | NA (NA-NA) | 0.55 | NA (NA-NA) | 0.53 | 0.9 (0.42-1.9) | 0.76 | 1.5 (0.94-2.6) | 0.1 |
|  | 1 | 0.075 (NA-3.5e+23) | 0.99 | 0.89 (0.67-1.1) | 0.35 | 0.96 (0.64-1.4) | 0.84 | 1.1 (0.95-1.2) | 0.22 |
|  | 2 | 1.1 (0.85-1.3) | 0.54 | 0.96 (0.8-1.2) | 0.64 | 0.89 (0.48-1.4) | 0.65 | 1 (0.95-1.2) | 0.37 |
|  | 3 | 1.1 (0.89-1.4) | 0.37 | 1 (0.83-1.3) | 0.87 | 0.68 (0.22-1.3) | 0.37 | 1.1 (0.96-1.2) | 0.27 |
|  | 4 | 1.1 (0.92-1.4) | 0.22 | 0.91 (0.76-1.1) | 0.35 | 0.73 (0.19-1.4) | 0.45 | 1.1 (0.96-1.2) | 0.24 |
| Cat IgG4 (µg/L)* | 6m | 0.057 (NA-2e+12) | 0.99 | 0.99 (0.67-1.3) | 0.95 | 24 (3.3e-30-NA) | 1 | 1.1 (0.88-1.3) | 0.41 |
|  | 1 | 0.76 (0.43-1.1) | 0.22 | 0.94 (0.78-1.1) | 0.54 | 0.76 (0.42-1.2) | 0.28 | 1 (0.9-1.1) | 0.82 |
|  | 2 | 1.4 (1.1-1.7) | **0.011** | 0.92 (0.67-1.3) | 0.59 | 0.96 (0.51-1.6) | 0.88 | 1.1 (1-1.3) | 0.053 |
|  | 3 | 1.3 (1-1.6) | 0.05 | 0.9 (0.63-1.4) | 0.59 | 0.86 (0.054-13) | 0.91 | 1.2 (1-1.4) | **0.033** |
|  | 4 | 1.4 (1.1-2) | **0.027** | 0.89 (0.64-1.3) | 0.49 | 0.54 (0.011-1.5) | 0.58 | 1.2 (1-1.5) | **0.034** |
| Couch grass IgG4 (µg/L)* | 6m | NA (NA-NA) | 0.55 | 0.062 (NA-1.3e+24) | 0.99 | 19 (2.5e-57-NA) | 1 | 1.3 (0.74-2.4) | 0.32 |
|  | 1 | 0.081 (NA-9.7e+23) | 0.99 | 1 (0.77-1.3) | 0.81 | 0.93 (0.6-1.4) | 0.71 | 1.1 (0.92-1.3) | 0.29 |
|  | 2 | 0.071 (NA-2.1e+22) | 0.99 | 0.88 (0.7-1.1) | 0.22 | 0.91 (0.61-1.3) | 0.61 | 1 (0.88-1.1) | 0.96 |
|  | 3 | 1.2 (0.99-1.6) | *0.061* | 0.85 (0.7-1) | 0.1 | 1.4 (0.88-2.2) | 0.16 | 1.1 (0.96-1.2) | 0.22 |
|  | 4 | 1.1 (0.91-1.4) | 0.28 | 0.72 (0.56-0.91) | **0.0074** | 0.88 (0.24-1.9) | 0.75 | 1 (0.91-1.2) | 0.69 |
| Phadiatop Infant IgG4 (PAU/L)* | 6m | 0.7 (0.45-0.91) | **0.03** | 1 (0.88-1.2) | 0.79 | 1.4 (0.96-2.4) | 0.12 | 0.98 (0.89-1.1) | 0.67 |
|  | 1 | 0.91 (0.72-1.2) | 0.4 | 0.73 (0.49-0.99) | *0.057* | 0.83 (0.29-1.5) | 0.64 | 0.93 (0.81-1.1) | 0.35 |
|  | 2 | 1.1 (0.89-1.3) | 0.49 | 0.97 (0.68-1.6) | 0.86 | 1.7 (0.93-7.7) | 0.2 | 1.1 (0.96-1.3) | 0.2 |
|  | 3 | 2.3 (1.1-6.8) | *0.091* | 0.23 (0.071-0.64) | **0.0076** | 1 (0.17-7.3) | 1 | 1.3 (0.96-1.8) | 0.16 |
|  | 4 | 1 (0.83-1.4) | 0.71 | 0.3 (0.097-0.85) | **0.028** | 0.42 (0.042-3.2) | 0.4 | 1.1 (0.88-1.3) | 0.61 |
| HDM IgG (mg/L)* | 1 | 25 (0.32-1.6E+04) | 0.19 | 3.3 (0.16-46) | 0.38 | 5.6E-03 (8.4E-06-0.57) | *0.058* | 2 (0.31-11) | 0.44 |
|  | 2 | 0.8 (0.15-3.5) | 0.78 | 0.97 (0.24-3.7) | 0.96 | 0.79 (0.031-18) | 0.88 | 1.3 (0.6-2.9) | 0.48 |
|  | 3 | 2.3 (0.14-35) | 0.54 | 0.48 (0.057-2.5) | 0.43 | 3.9 (0.26-96) | 0.34 | 2.1 (0.89-5) | *0.089* |
| Cat IgG (mg/L)* | 1 | 1.5E-15 (NA-1.2E+291) | 0.99 | 6.5 (0.22-150) | 0.24 | 4.6E-03 (1.4E-06-0.9) | *0.082* | 1.7 (0.11-18) | 0.68 |
|  | 2 | 0.66 (0.077-3.5) | 0.65 | 1.2 (0.28-4.3) | 0.82 | 0.16 (4.0E-03-3.5) | 0.26 | 0.87 (0.34-2.1) | 0.75 |
|  | 3 | 0.023 (8.2E-06-2) | 0.18 | 0.52 (0.058-2.7) | 0.49 | 3.7 (0.18-244) | 0.44 | 1.1 (0.35-3) | 0.9 |

BMI = body mass index; HDM = house dust mite; LRI = lower respiratory infection. Association analyses performed via generalised linear models (GLM) with demographic covariates: age-five wheeze ~ predictor + sex (male) + BMI at age 3 + paternal history of asthma + maternal history of asthma. Bold text indicates statistical significance ( $p < 0.05$ ); italics indicate near-significance ( $p < 0.10$ ). \*Odds ratio (OR) is for every 10-fold increase in IgE, IgG4 or IgG.

### 2.2 CAS2: low-risk cluster susceptible to atopic and non-atopic wheeze

Like CAS1, CAS2 was a low-risk cluster with infrequent allergic disease. However, compared to CAS1, Phadiatop and HDM IgE were slightly elevated at most timepoints (**Table 2, Supplementary Table 3B**; Figure 3A). Conversely, peanut IgE was not significantly elevated (Wilcoxon, adjusted *p*=0.99 at all timepoints; **Figure 3D**). As for other antibody isotypes: CAS2 IgG was between CAS1 and CAS3 levels, with it being closer to CAS1. CAS2 IgG4 was also intermediate, but was much closer to CAS3 levels than CAS1 (**Table 2; Figure 3**). However, despite the antibody differences between CAS1 and CAS2, yearly rates of wheeze in CAS2 remained comparable to CAS1 (30% at age 1, declining to 18% at age 10; **Table 1; Figure 2**). Interestingly, compared to CAS1, individuals in CAS2 tended to have fewer older siblings living in the household at age 2, as well as more frequent paternal history of asthma (adjusted *p*=0.029 and 0.055, respectively; **Table 1**).

**Figure 3:**
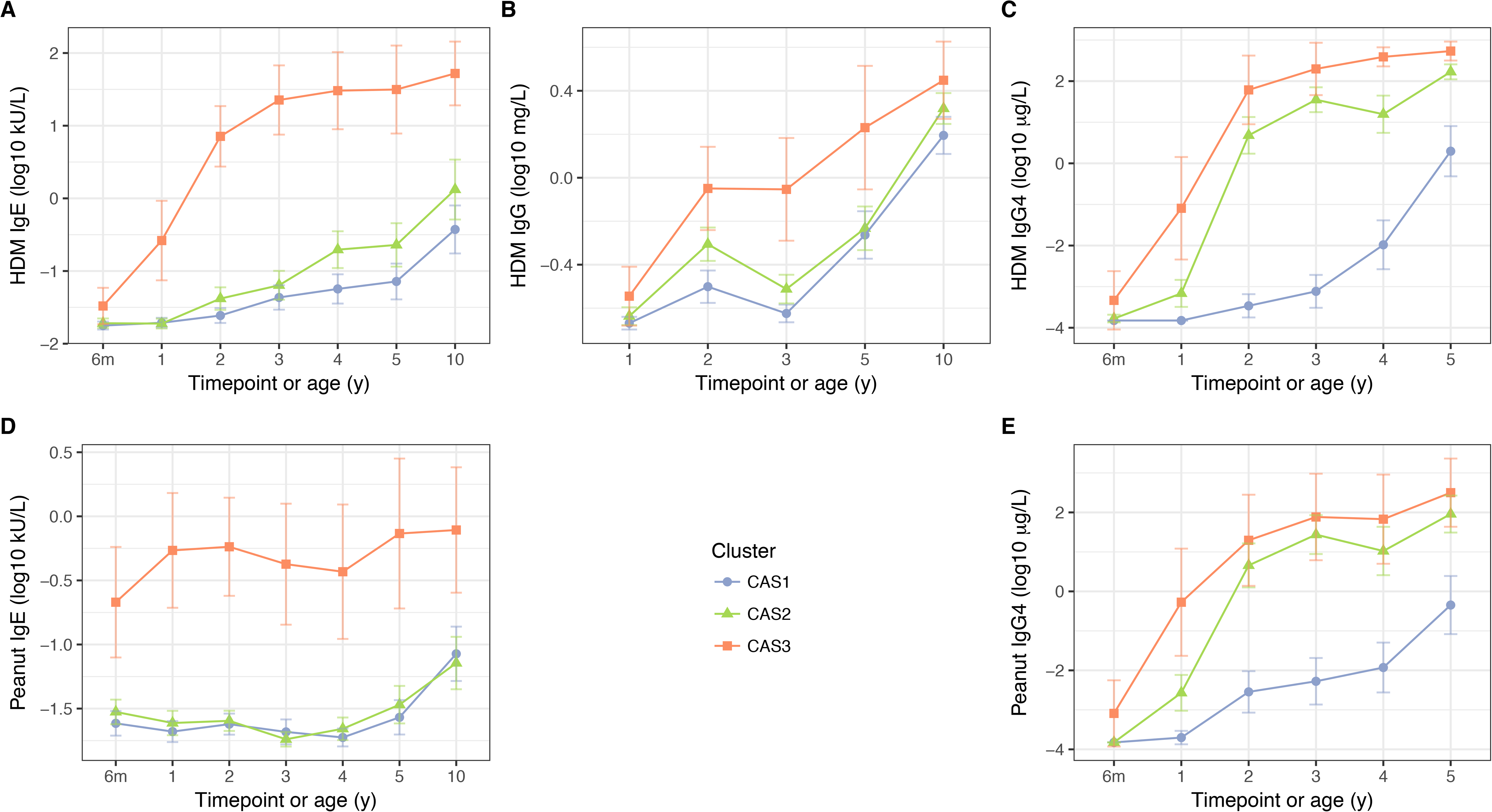
HDM IgE (A), IgG (B) and IgG4 (C); and peanut IgE (D) and IgG4 (E) stratified by cluster and time, in the CAS dataset. Points indicate means; bars indicate 95% CI (t-distribution).

The predictive factors for wheeze at age 5 in CAS2 included: LRI, wLRI and fLRI frequency (GLM; *p*=2.7×10^−3^, 0.016 and 0.02 at age 3, respectively); HDM IgE (*p*=0.016 and 0.011 at ages 2 and 4, respectively); and Phadiatop IgE (*p*=0.01 at age 4) (Table 4; Figure 4). After multiple regression analysis with stepwise backward elimination (Methods), three of these remained significant: age-two fLRI (*p*=0.006, OR 11 per unit increase), age-four wLRI (*p*=0.006, OR 4.8), and age-four Phadiatop IgE (*p*=5.5×10^−3^, OR 4.1). Repeated-measures ANOVA showed that HDM IgE and LRI-related variables (LRI, wLRI, fLRI) from the first 3 years of life were significant predictors of age-five wheeze in CAS2 (**Supplementary Table 4**). But although both allergic (IgE-related) and non-allergic (infection-related) risk factors contributed to age-five wheeze, there was no significant evidence of interaction between them (*p*=0.36 within CAS2 alone, *p*=0.92 across entire cohort, for age-four wLRI frequency × Phadiatop IgE). Overall, CAS2 represented a low-risk trajectory susceptible to, but not necessarily afflicted by, wheeze due to atopic and non-atopic risk factors. In this cluster, atopic determinants of age-five wheeze (HDM and Phadiatop IgE) were only active from age 2 onwards, suggesting delayed atopic wheeze in this cluster.

**Figure 4:**
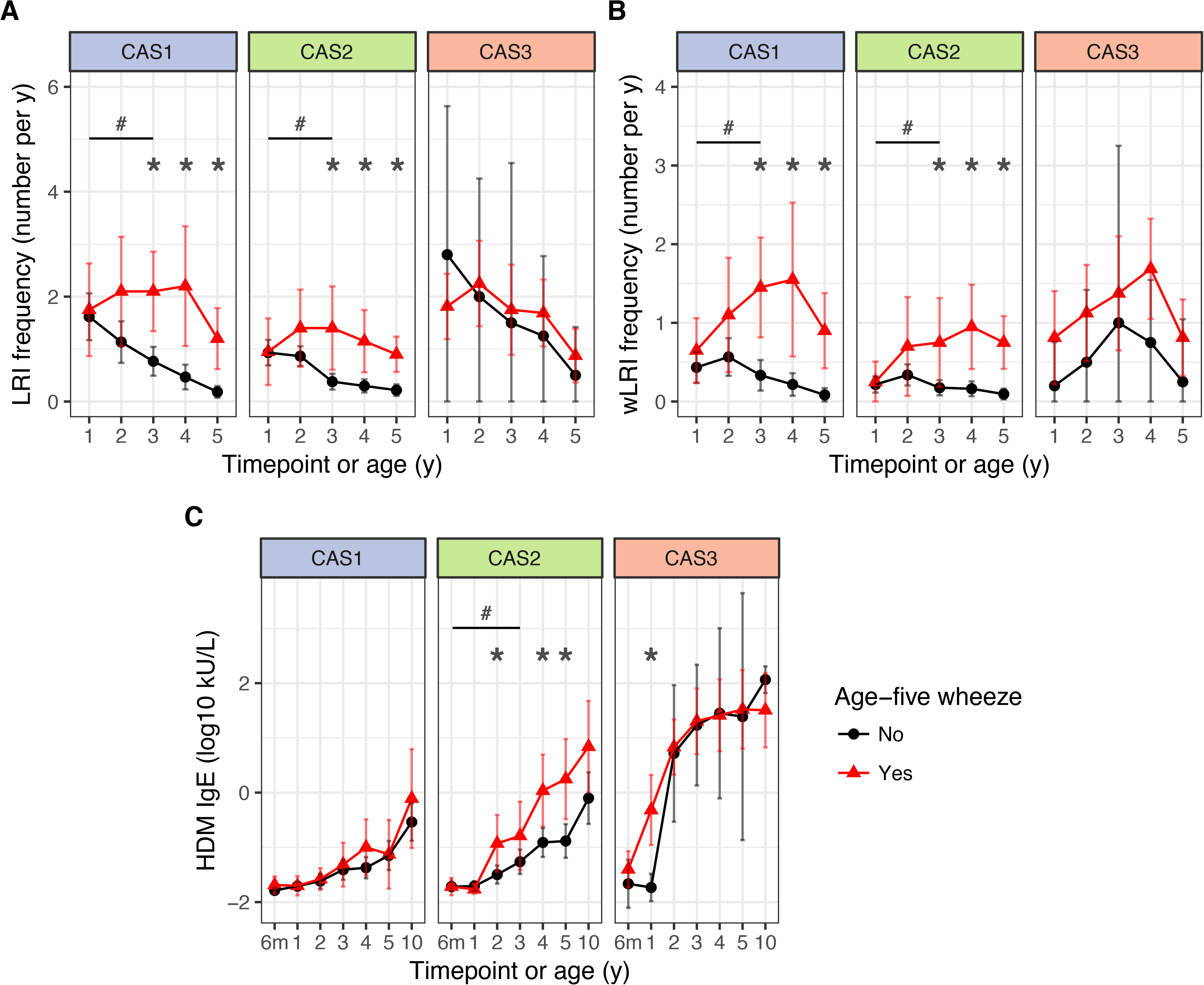
LRI frequency (A), wheezy LRI (wLRI) frequency (B), and HDM IgE (C), stratified by age-five wheeze status, cluster and time, in the CAS dataset. Points indicate means; bars indicate 95% CI (t-distribution). \**p*<0.05 for Mann-Whitney-Wilcoxon comparison within each timepoint. #*p*<0.05 for repeated-measures ANOVA across timepoints from the first 3 years of life (see Table 4).

### 2.3 CAS3: high-risk atopic cluster with persistent wheeze

CAS3 was a “high-risk” cluster, where persistent respiratory wheeze and atopic disease was seen in more than half the group throughout the first 10 years of life (**Table 1; Figure 2**). This cluster was dominated by males (86%, Fisher exact test, unadjusted *p*=6.8× 10^−3^ compared to CAS1, **Table 1**), and appeared to represent an early- and multi-sensitised atopic phenotype with persistent wheeze.

CAS3 had elevated IgE, IgG, and IgG4 responses to common allergens, especially HDM (**Table 2, Supplementary Table 3B**; Figure 3). Peanut, HDM and Phadiatop IgE were significantly greater in CAS3 than in CAS1 from 6 months onwards. SPTs were also more frequently positive in CAS3, especially to HDM and food allergens. From 6 months onwards, wheal sizes in cow’s milk and egg white SPTs were on average greater in CAS3 than in CAS1 (Wilcoxon, adjusted *p*=4.8×10^−9^ for egg white SPT at age 5, **Supplementary Table 3D**). Age-five peanut SPTs also yielded stronger wheal responses in CAS3 compared to CAS1 (Wilcoxon, adjusted *p*=8.4×10^−4^).

No strong predictors for age-five wheeze were identified within CAS3 (**Table 4**): only couch grass-specific IgE at age 2 and ARI frequency at age 1 were weakly significant (both *p*=0.046). Neither of these reached statistical significance with stepwise backward elimination. However, the prolific IgE response, and the prevalence and severity of early-life LRIs in this cluster (**Table 3**), strongly suggest contribution from both atopic and non-atopic causes of wheeze. CAS3 primarily represented those with extreme levels of atopic sensitisation and infection. The relative paucity of identifiable predictors may be explained by the small size of CAS3 (*N*=22), the intrinsically high rate of wheeze in the cluster (76% with age-five wheeze), and saturation of risk from high levels of IgE and frequent infections.

### 2.4 Cytokine responses of PBMCs following in vitro antigen stimulation

Unlike the antibody measurements, no cytokine measurements contributed as clustering features to the original cluster analysis. Nonetheless, we found that *in vitro* stimulation of PBMCs with HDM antigen elicited a stronger Th2 cytokine response in CAS3 compared to the other clusters (**Table 2, Figure 5**). These cytokines (IL-4, IL-5, IL-13) were elevated from a very young age (Wilcoxon, adjusted *p*=4.6× 10^−5^ for IL-4 mRNA at age 6m, compared to CAS1), coinciding with increase in HDM IgE and IgG4 responses. Similar differences in CAS3 were observed for peanut- and ovalbumin-stimulated PBMCs, but only at 6 months of age (unadjusted p<0.05 for all, **Supplementary Table 3C**). There were no other significant differences for other non-Th2 cytokines that were tested (IFN-γ, IL-10), nor were there cytokine differences specific for CAS1 or CAS2 (**Supplementary Table 3C**).

**Figure 5:**
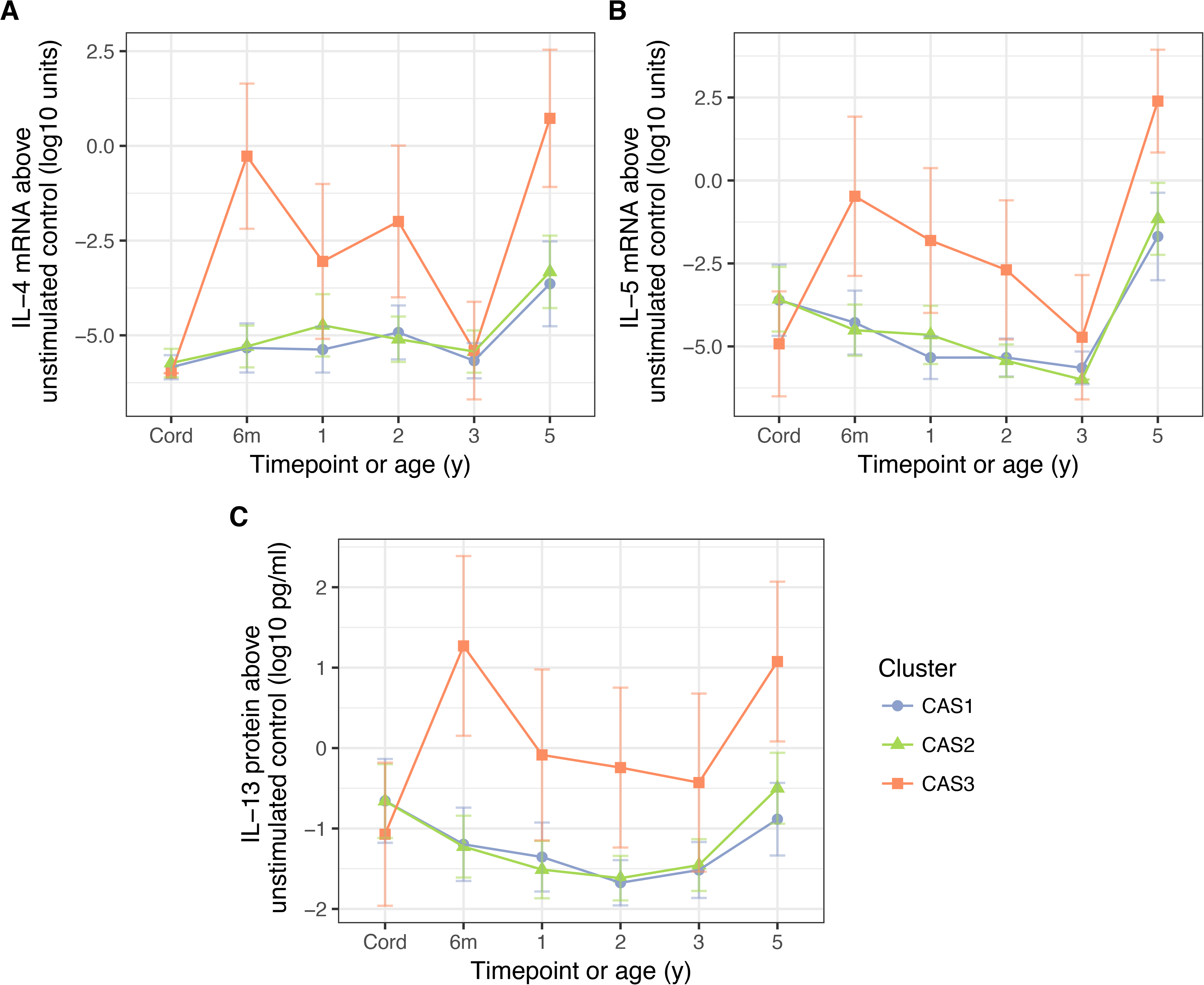
PBMC expression of IL-5 (A) and IL-4 mRNA (B), as well as IL-13 protein (C), in response to stimulation HDM, stratified by cluster and time (CAS) Cord = cord blood sample collected at birth. Points indicate means; bars indicate 95% CI (t-distribution).

### 2.5 IgG4 and IgG

Across all clusters, allergen-specific IgG4 and IgG were positively correlated with IgE for the same allergen (especially HDM, **Supplementary Figure 2**). As noted previously, CAS2 and CAS3 were distinguished from CAS1 by high IgG4 against multiple allergens, and CAS3 had greater IgG4 responses than either CAS1 or CAS2 (**Supplementary Table 3B**; Figure 3). Although previous literature suggests possible protection conferred by IgG4 [28] or IgG [29], in this study there was no clear or consistent evidence of protection by either IgG4 or IgG against later wheeze (**Table 4**). Furthermore, the protected status of CAS2 was unlikely to be driven by IgG4, given that CAS3 had higher IgG4 than CAS2.

### 2.6 Patterns in IgE, IgG, cytokine and SPT responses

Although they were highly-correlated, phenotypes of IgE, IgG, Th2 cytokine and SPT responses did not overlap perfectly. CAS3 was enriched for individuals with strong signals in all modalities, but there remained individuals within CAS3 and the rest of the cohort who were only responsive in some modalities but not others. Notably, IgE and SPT signals did not always coincide (**Supplementary Figure 3A**). Also, some individuals with IgE or SPT sensitisation against HDM did not exhibit detectable Th2 cytokine response to *in vitro* HDM stimulation (**Supplementary Figure 3A**). Finally, HDM IgG4 did not appear to be responsible for this effect: those with IgE but not Th2 cytokine responses (i.e. HDM IL-13 at limit of detection 0.01 pg/L) did not have significantly different levels of IgG4 compared to those with both IgE and Th2 responses (Wilcoxon, *p*=0.15, **Supplementary Figure 3B**).

### 2.7 Comparison to existing criteria for atopy

The information conveyed by the npEM-derived CAS clusters was consistent with that of traditional atopy thresholds (i.e. any specific IgE ≥ 0.35 kU/L or SPT ≥ 2mm at age 2). When we compared the CAS clusters with supervised groups created using traditional thresholds (**Supplementary Table 5**), we found that CAS1 most closely matched a non-atopic phenotype (58 of 84 had no specific IgE greater than 0.35 kU/L by age 2). Conversely, CAS2 and CAS3 partially matched the traditional criteria for atopy, with CAS3 being an extreme phenotype (all 22 children in CAS3 had some specific IgE ≥ 0.35kU/L by age 2).

However, the CAS clusters outperformed IgE- and SPT-defined atopy in terms of predicting for age-five wheeze (likelihood ratio test for model with clusters vs. model with IgE/SPT, Chi-squared=23, *p*=2.0×10^−6^). In addition, at age 2, 68% of CAS3 were “sensitised” (any specific IgE ≥ 0.35kU/L) to two or more allergens, compared to only 1% and 6% for CAS1 and CAS2 respectively. This emphasised CAS3 as an early- and multi-sensitised phenotype. Finally, many members of low-risk CAS1 and CAS2 who were IgE- or SPT-responsive prior to age 5 did not maintain atopic wheeze at age 5 (77% or 79 of 103), compared to CAS3 (24% or 5 of 21). Therefore, the association of IgE and SPT results with disease risk varied across clusters. Overall, this suggests that fixed atopy thresholds are not sufficient on their own in delineating risk profiles – instead, an unsupervised clustering approach may be superior.

### 2.8 Relationship with time-dependent wheeze phenotypes

We explored how the npEM-derived clusters mapped to pre-defined wheezing phenotypes (**Figure 2C**): no wheeze (in the first three years of life, or at age 5), transient wheeze (only in the first three years of life), late wheeze (only at age 5), and persistent wheeze (in both first three years of life and age 5). We found that CAS3 was enriched for persistent wheeze, while individuals in CAS1 or CAS2 tended to have transient or no wheeze. There were rarely any members of the cohort with late wheeze (approximately 10% or less).

### 2.9 Co-associations with food sensitisation, eczema and wheeze

In addition to persistent wheeze, CAS3 was also enriched for persistent food sensitisation (peanut IgE ≥ 0.35 kU/L, or positive egg white or cow’s milk SPTs) and persistent eczema: 44% of all individuals in CAS3 satisfied all three conditions (**Supplementary Figure 4**). Almost all individuals in CAS3 had both eczema and food sensitisation from age 6m onwards, with rates of food sensitisation and wheeze increasing with time (**Figure 2D**). In contrast, CAS1 and CAS2 had low rates of food sensitisation, and declining rates of both eczema and wheeze. These trends lend credence to the hypothesis that the “atopic march” phenotype [30, 31] may only be present in a minority of the population (e.g. CAS3) [19].

### 2.10 Relationship with microbiome

Previous studies suggest an association between asthma risk and early-life disruption of the respiratory microbiome, especially colonisation with *Streptococcus* spp. in the first 7 weeks of life [24]. In this study, using the same data and definitions, we found that CAS3 was overrepresented by individuals who had >20% relative abundance of *Streptococcus* in their first infection-naïve healthy NPA, within the first 7 weeks of life (44% versus 11% and 15% in CAS1 and CAS2, respectively; Fisher exact test, unadjusted *p*=0.042 and 0.065, respectively; **Table 3**).

Furthermore, Teo et al and others [24, 32] previously found that transient incursions with MPGs associated with acute respiratory infections (*Streptococcus, Haemophilus* and *Moraxella* spp.) were associated with increased frequency and severity of subsequent LRIs and wheezing disease. Here, we found that the proportion of infection-associated MPGs in healthy samples from age 0 to 2 was greater in CAS3 (62% vs. 49% and 32% in CAS1 and CAS2, respectively; Fisher exact test, unadjusted *p*=0.2 and 5.5×10^−4^, respectively; **Table 3**). This finding was independent of LRI and wLRI frequency (GLM; *p*<0.05 for model predicting group membership, with age-two LRI and wLRI as covariates). On the contrary, there were no associations between cluster membership and health-associated MPGs *(Corynebacterium, Alloiococcus, Staphylococcus* spp.; **Supplementary Table 3E**).

Recent work by Teo et al [25] suggested that infection-associated MPGs in early life were predictive for age-five wheeze in atopic children, while in non-atopic children they were predictive for transient wheeze (i.e. wheeze only in the first 3 years of life and not later). In this study, a similar trend was noted for infection-associated MPGs from age 0 to 2, in relation to transient wheeze in “non-atopic” CAS1 (GLM, OR 3.6 per percent, *p*=0.17, with demographic covariates). Surprisingly, there was evidence that infection-associated MPGs in later samples (from age 2 to 4) were *protective* against age-five wheeze in CAS1 (OR 0.086 per percent, 0.45 per quartile, *p*=0.034 and 0.035, respectively; **Table 4**). Infection- and health-associated MPGs were otherwise not associated with age-five wheeze within the other clusters.

### 2.11 Decision tree analysis

We used decision tree analysis to determine the handful of biological features that most strongly distinguish each npEM cluster. This process may allow us to simplify the clustering into a tree algorithm that can then be used clinically for screening or diagnosis. Unlike the previous GLMs, which identified variables most predictive for wheeze, the decision trees identified variables most discriminatory for age-five wheeze versus non-wheeze within each cluster.

Decision tree analysis on the CAS dataset, using all available variables from all timepoints for classification, created a “Simple Tree” with two decision nodes and three end nodes (**Figure 6**). This tree had 89% accuracy in terms of retrieving the cluster memberships from the original npEM model, where accuracy is calculated as percentage overlap of tree clusters with original CAS clusters. Applying this Simple Tree to CAS, we found that membership in the CAS3-equivalent tree cluster was also a better predictor for age-five wheeze (likelihood ratio test, Chi-squared=19, p<1×10^−5^) than traditional thresholds for atopy based on IgE and SPT measurements at age 2.

**Figure 6:**
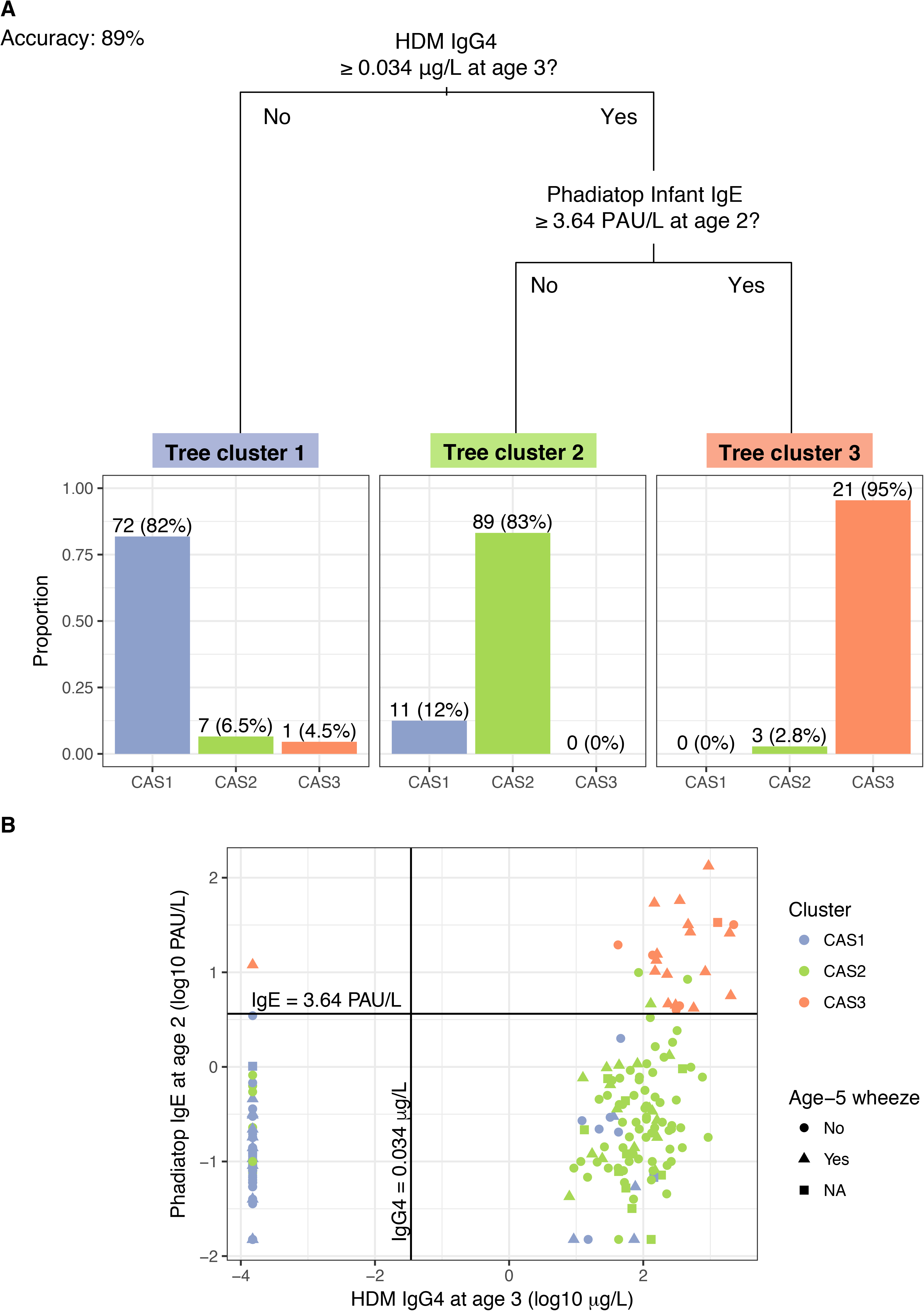
A “simple” decision tree generated by recursive partitioning from CAS data, with breakdown of tree clusters by actual CAS npEM-derived clusters (A); scatterplot showing separation of CAS clusters by decision split thresholds (B) Percentages in Panel A may not sum up to 100%, because some individuals have missing values for decision node variables, hence making them impossible to classify. In Panel B, note that left-most column of points represent values of HDM IgG4 that were less than the limit-of-detection (LOD) for that assay (0.0003 μg/L), and were subsequently assigned to half the LOD (0.00015 μg/L). Most of these points belonged to individuals from CAS1.

Further tree analyses using variables restricted to each timepoint identified similar trends (**Supplementary Figure 5**); IgG4-related variables best separated CAS1 from other clusters (from Phadiatop in early life, to HDM by age 3), while IgE-related variables (Phadiatop) best separated CAS2 and CAS3. Explicitly forcing the exclusion of Phadiatop variables from tree analysis caused these thresholds to be replaced with allergen-specific assays: cat and peanut IgG4 for Phadiatop IgG4; and peanut and HDM IgE for Phadiatop IgE (**Supplementary Figures 6 and 7**). This is consistent with correlation patterns amongst the IgE and IgG4 variables (**Supplementary Table 6**).

Because the causes of wheeze were likely different for different clusters, we also constructed a “Comprehensive Tree” that best split individuals into six groups, based on cluster membership crossed with age-five wheeze status (**Supplementary Figure 8**). For decision nodes we excluded all age-five features related to wheeze (e.g. LRIs, wheezy LRIs at age 5), because of definitional overlap with our outcome of interest. We thus identified nodes that were consistent with the predictors for wheeze found in the previous regression analyses (**Table 4**), combined with nodes from the Simple Tree (**Figure 6**). The Comprehensive Tree had a total accuracy of 77% in correctly recovering both cluster membership and wheeze status. In terms of identifying purely wheeze status at age 5, the accuracy of the tree was 84%, with a positive predictive value (PPV, or precision) of 72%, negative predictive value (NPV) of 88%, sensitivity (recall) of 71% and specificity of 89%. The Comprehensive Tree was more successful in flagging age-five wheeze (likelihood ratio test, Chi-squared=60, *p*=6.1 × 10^−13^), compared to the traditional atopy thresholds described previously.

### 2.12 External replication of clusters in MAAS and COAST

The disease trajectories described by the CAS npEM clusters were successfully replicated in both MAAS (N=1085) [26] and COAST (N=289) [27]. We applied our npEM classifier (**Methods**) to MAAS and COAST, and found that individuals classified into “Cluster 3” (MAAS3/COAST3) had a persistent disease phenotype extending into late adolescence, with consistently high rates of parent-reported wheeze and physician-diagnosed asthma from birth to age 16. The other two clusters (Cluster 1 = MAAS1/COAST1; Cluster 2 = MAAS2/COAST2) appeared to be relatively low-risk (**Figure 7A,B,D**).

**Figure 7:**
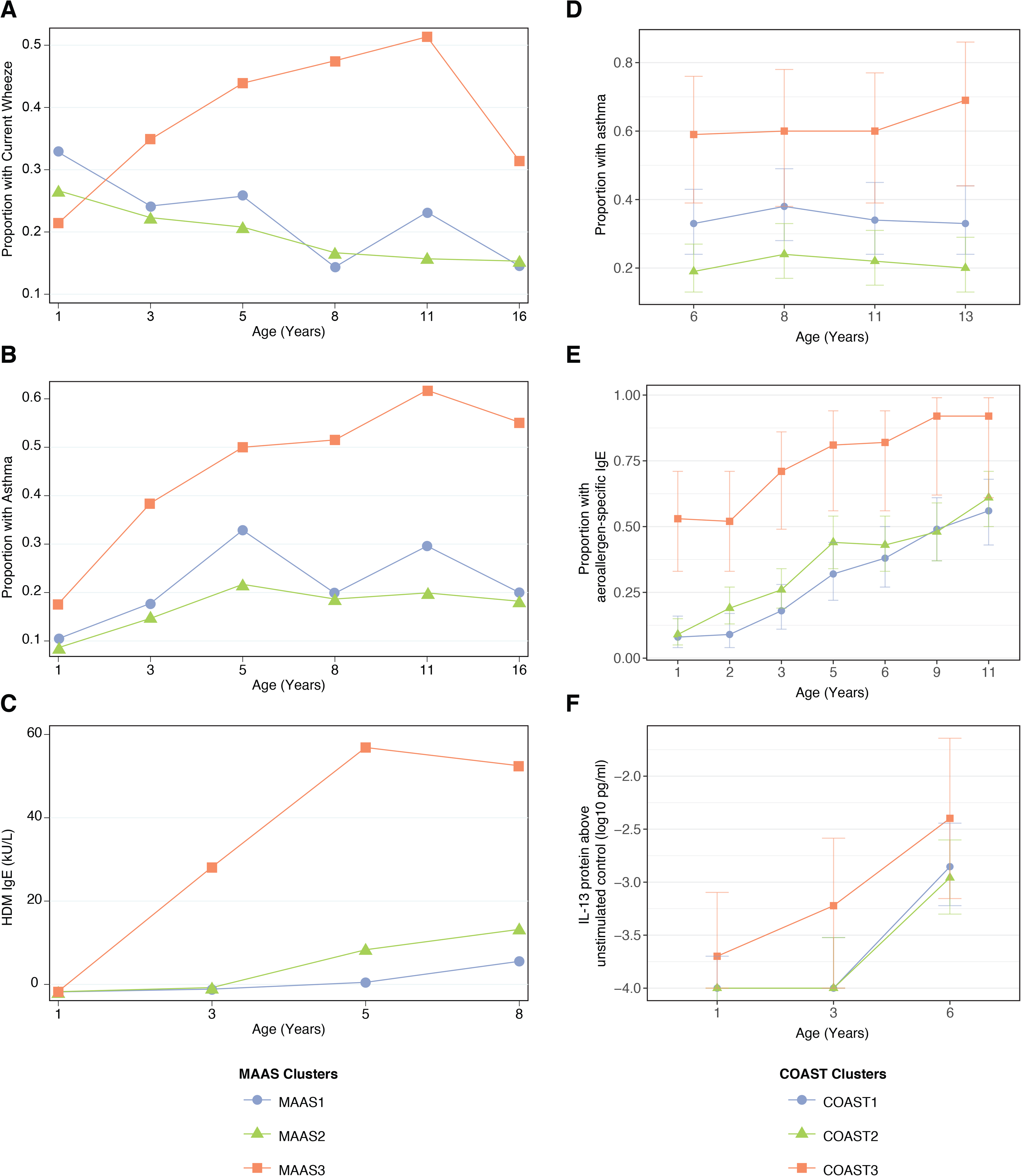
Description of npEM-derived clusters in external cohorts: in MAAS, incidence of wheeze (A), asthma diagnosis (B), and HDM IgE levels (C); in COAST, incidence of asthma diagnosis (D), proportion of individuals with detectable aeroallergen-specific IgE levels (E), and PBMC protein expression of IL-13 following HDM stimulation above unstimulated control (F) MAAS cohort (N=934) was classified using npEM model from CAS, into MAAS1 (N=199, 21%), MAAS2 (N=692, 74%) and MAAS3 (N=43, 5%); these correspond to CAS clusters CAS1, 2 and 3, respectively. COAST cohort (N=285) was similarly classified into COAST1 (N=105, 37%), COAST2 (N=151, 53%) and COAST3 (N=29, 10%).

MAAS3 and COAST3 exhibited stronger IgE expression (total, HDM, cat, dog) from ages 1 to 8 (**Figure 7C,E**), compared to other clusters in each dataset. Like CAS3, COAST3 demonstrated elevated PBMC expression of Th2 cytokine protein (IL-5 and IL-13) in response to HDM stimulation at age 3 (**Figure 7F**). This was not replicated in MAAS3, but previous work in MAAS had identified that a strong Th2 response (IL-5, IL-13) to HDM stimulation of PBMCs at age eight was associated with increased risk of HDM sensitisation and asthma [21]. Nonetheless, MAAS3 appeared to be overrepresented in “early-sensitised” and “multiple sensitised” phenotypes discovered earlier by Lazic et al [17] from SPT and IgE data. Approximately 86% of individuals in MAAS3 belonged to either one of these two phenotypes, although only 13% of individuals in these two phenotypes were accounted for by MAAS3.

**Figure 8:**
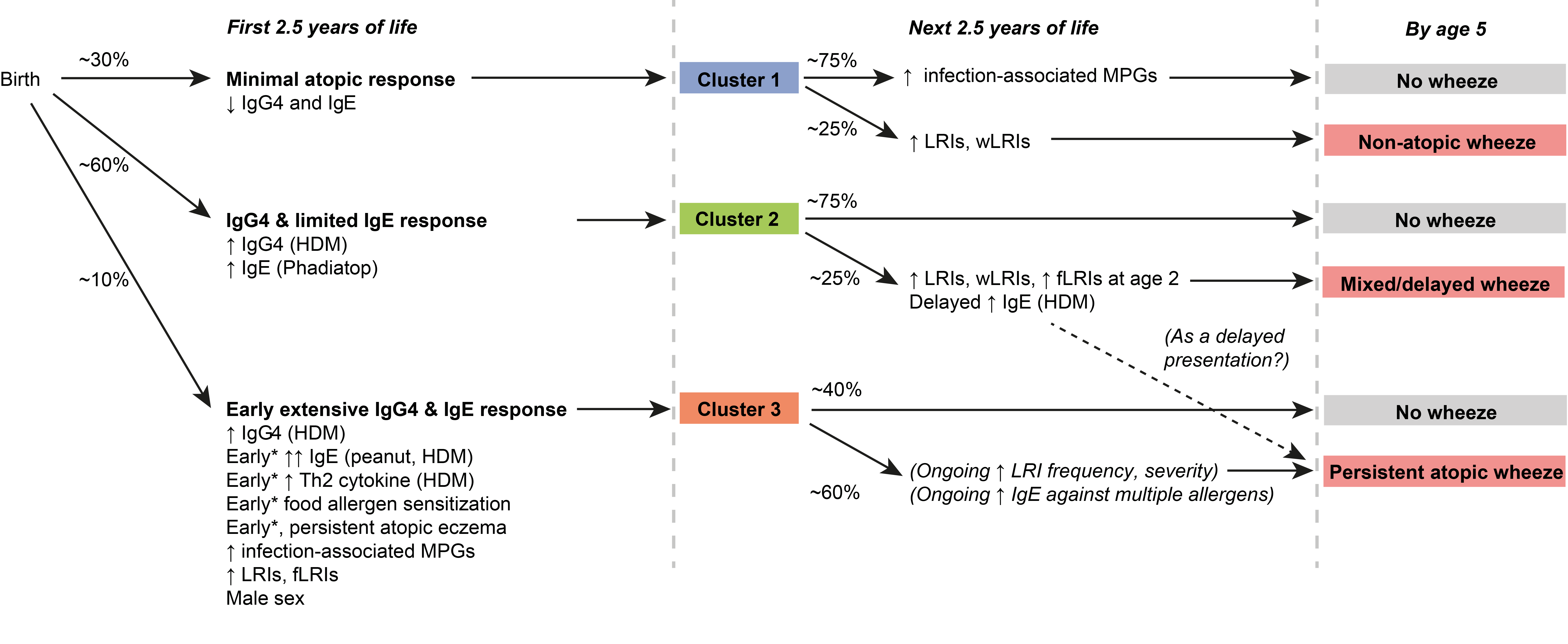
Graphical summary of proposed clusters. *”Early” specifically refers to “within the first 6 months of life”.

Furthermore, when we explored potential predictors of wheeze phenotypes and asthma diagnosis in later childhood, we found that the clusters in the external cohorts were very similar to those in CAS. In COAST1, LRI and wLRI frequency at age 2 were predictive of asthma diagnosis at age six (GLMs with demographic covariates, *p*=0.02 and 0.02, respectively), while in COAST2, HDM IgE at age 3, and LRI, wLRI and fLRI frequencies at age 2 were all predictive (GLMs, *p*<0.05 for all) (**Supplementary Figure 9**). Although the timing and magnitude of associations differed between cohorts, this reaffirmed wheeze in Cluster 1 as being primarily non-atopic in origin, while wheeze in Cluster 2 seemed to be driven by both non-atopic and atopic factors.

We attempted to validate CAS-derived decision trees in the MAAS dataset, as it contained measurements of both age-five HDM IgE and HDM IgG4, which we used as surrogates for age-three HDM IgE and HDM IgG4. These features comprised two decision-node features in the Phadiatop-free equivalent of the CAS Simple Tree (**Supplementary Figure 5**). COAST did not have any IgG4 measurements, so tree validation was not attempted there. The performance of the Simple Tree when applied to MAAS was poor, with only 20% accuracy in terms of overlap between tree clusters and npEM clusters, compared to 89% in CAS. Instead, when we generated a new tree from MAAS using its npEM clusters (**Supplementary Figure 10**), we achieved good accuracy (85% clusters correctly identified). However, the decision nodes of this tree were different to CAS, being related to family size and SPTs rather than IgE and IgG4. The MAAS1-equivalent tree cluster was distinguished by reduced SPT responsiveness (wheal size < 4-5mm) to cat, HDM and grass; while the MAAS3 equivalent was defined by strong HDM or grass SPT responsiveness. While this differs from the CAS decision trees, it is consistent with the broad distinction between non-atopic Cluster 1 and atopic Cluster 3. We also stress that CAS and COAST are both high-risk cohorts (each child having a parent with asthma or allergies), while MAAS was not.

## 3 Discussion

We have used model-based cluster analysis to uncover clusters of children with differential asthma susceptibility. Specifically, there was a high-risk group characterised by very early allergen-specific Th2 activity; early sensitization to multiple allergens including food allergens; and concurrent frequent respiratory infections – resulting in high incidence of atopic persistent wheeze. We also found a lower-risk cluster, with limited or delayed elevation in IgE – this resulted in a lower incidence of mixed (atopic and non-atopic) wheeze. Finally, there was a low-risk cluster which exhibited occasional and transient infection-related wheeze, with minimal allergen sensitisation. These clusters were replicated in external datasets, suggesting relevance across populations. A decision tree that accounts for cluster membership with modified thresholds for atopic sensitisation is superior to traditional definitions of atopy in predicting disease occurrence. A summary of key findings is given in **Table 5**, and in-depth discussion of the biological and practical significance of these findings can be found in the **Supplementary discussion**.

**Table 5:** Key findings from cluster analysis

|  |
| --- |
| ▪ Certain childhood populations may be broadly split into three clusters, each representing a unique trajectory of immune function and susceptibility to respiratory infections: low-risk non-atopic Cluster 1 with transient wheeze; low-risk but allergy-susceptible Cluster 2 with mixed wheeze; and strongly-atopic high-risk Cluster 3 with persistent wheeze.<br>▪ Cluster 3 is consistent with an early-sensitised and multi-sensitised phenotype.<br>▪ HDM hypersensitivity is an important predictor of wheeze in allergic or allergy -susceptible individuals.<br>▪ In CAS, IgG4 flags for clusters with susceptibility to atopic disease (CAS2 and CAS3), while early and multiple-allergen elevation in IgE predicts frank atopic disease. The pathophysiological role of IgG4 remains unclear.<br>▪ Food and peanut hypersensitivities are important contributors to membership in high-risk Cluster 3. This may be pathophysiologically related to eczema, multi-sensitisation and the atopic march.<br>▪ Allergic and infective processes act in a synergistic manner to intensify airway inflammation during respiratory pathogen clearance. Some clusters (Cluster 3) may be more susceptible to this effect than others that lack strong allergic sensitisation (Cluster 1).<br>▪ The microbiome also acts differently on asthma risk depending on cluster membership. In CAS, early-life asymptomatic colonisation with infection-associated MPGs is associated with risk of persistent wheeze in allergy-susceptible clusters (CAS2, CAS3), while it is potentially protective in non-atopic children (CAS1)<br>▪ Tests for atopy (IgE, SPT, cytokines) do not necessarily overlap. Therefore, atopy may be better defined by the composite result from a battery of tests encapsulated in a predictive model, rather than just a single test or threshold.<br>▪ Different childhood populations may share similar trajectories of asthma susceptibility, but there may be subtle differences in terms of the types of tests, allergens, or biological signals that are most informative (SPT, IgE, cytokines, etc.). |

Our findings demonstrate clear and homogeneous developmental trajectories among children in multiple cohorts. The latent clusters incorporated multiple domains, including immune function and infection frequency, that reflected both endotype (pathophysiology) and phenotype. We emphasise that our approach was unsupervised and exploratory – endpoint variables describing parent-reported wheeze and atopic disease were excluded from clustering, and clusters were constructed *de novo* without any reference to traditional atopic thresholds. Therefore, it was encouraging to observe that the data-driven clusters differed in susceptibility and nature of subsequent wheeze, and that biologically- and clinically-relevant findings could still be derived from them. Our results build on previous findings [11, 33] demonstrating that the concept of atopy, as an intrinsic or heritable predisposition to allergic disease, is more complicated than what could be described by dichotomies or thresholds. Instead, our study strongly supports the future use of predictive models with more precise, subgroup-driven representations of atopy and other relevant pathophysiological mechanisms.

The characterisation of these three clusters demonstrates how the complex phenomenon of asthma pathogenesis can be explored in depth using clustering. We have successfully provided an example where addressing inter-cluster differences have allowed the identification of intra-cluster disease predictors. The clusters may be further characterised by exploring other aspects of asthma pathophysiology, including genetics, epigenetics and others. By continuing with these approaches, we can hopefully move away from fixed thresholds or criteria for atopic risk, to more sophisticated formulations of risk, which will then improve future attempts at the targeted screening, prevention and treatment of asthma. These approaches may also be broadly applied to other heterogeneous diseases or datasets, and computerised tools may then be designed to embody the sum knowledge from these approaches. Such approaches can eventually help clinicians and scientists achieve a fuller understanding of pathophysiology, and hence better predict and manage human disease.

## 4 Methods

### 4.1 Patients and study design

The Childhood Asthma Study (CAS) was a prospective birth cohort (*N*=263) operated by the Telethon Kids Institute, Perth, Western Australia [22]. CAS was established with the goal of describing the risk factors and pathogenesis of childhood allergy and asthma. Details of CAS have been reported in previous publications from our group [22, 24, 34–36], and are summarised below.

In CAS, expectant parents were recruited from private paediatric clinics in Perth during the period spanning July 1996 to June 1998. Each child who was born and subsequently recruited had at least one parent with physician-diagnosed asthma or atopic disease (hayfever, eczema). The child was then followed from birth till age 10 at the latest, with routine medical examinations, clinical questionnaires, blood sampling at multiple time points (6-7 weeks, 6 months, 1 year, 2, 3, 4, 5, and 10 years) and collection of nasopharyngeal samples. Parents also kept a daily symptom diary and recorded the presence of symptoms of respiratory infection and other illnesses, and all medications taken.

### 4.2 Measurements

During each routine visit, we collected samples that encompassed multiple domains of childhood health, and recorded metrics related to suspected or known modulators of asthma risk. These included markers of immune function, specifically: 1) IgG, IgG4, and IgE Phadiatop ImmunoCAP antibodies (ThermoFisher, Uppsala, Sweden), covering common allergens such as house-dust mite (HDM, *Dermatophagoides pteronyssinus)*, mould, couch grass, ryegrass, peanut, cat dander; 2) IgE and IgG4 Phadiatop Infant and Adult assays (ThermoFisher, Uppsala, Sweden) that target multiple allergens simultaneously [23]; 3) skin prick or sensitisation tests (SPT), testing for HDM, mould, ryegrass, cat, peanut, cow’s milk and hen’s egg; and 4) cytokine responses (IL-4,5,9,13,10, IFN-γ) following in vitro stimulation of extracted peripheral blood mononuclear cells (PBMCs) by multiple antigen and allergen stimuli, including phytohaemaglutinin (PHA), HDM, cat, peanut and ovalbumin. Additional details on these measurements are found in Hollams et al [34] and Holt et al [35].

In addition, nasopharyngeal samples were taken from each child during healthy routine visits, as well as unscheduled visits where parents were asked to present with their child at every onset of symptoms of a respiratory infection. We then screened these samples for viral and bacterial pathogens using rtPCR and 16s rRNA amplicon sequencing with Illumina MiSeq (San Diego, US), respectively [24]. Specific details are described in the supplement to Teo et al [24].

Other collected data included: sex, height and weight; paternal and maternal history of atopic disease; blood levels of basophils, plasmacytoid and myeloid dendritic cells as measured by fluorescence-assisted cell sorting (FACS); and levels of vitamin D (25-hydroxycholecalciferol, 25(OH)D), the measurement of which has been described by Hollams et al [36].

The study designs and measurements performed in the replication cohorts (MAAS, COAST) have been described elsewhere [19, 37]. Respiratory infection phenotypes (ARI, LRI, URI, fLRI, wLRI) were redefined in COAST based on their recorded symptom scores.

### 4.3 Identification of latent clusters

We used an implementation of a non-parametric mixture model (npEM) from the R package “mixtools” [38], because: 1) it was plausible to consider a population as a mixture of subpopulations each with their own unique distributions; 2) it had advantages over other unsupervised approaches [39] – unlike LCA, npEM could handle continuous variables; and 3) it lent itself to an intuitive method for supervised classification of other datasets into similar clusters (see **Supplementary Methods**). Further details can be found in the supplementary, and a graphical outline of methodology is given in **Supplementary Figure 1**.

Prior to cluster analysis, quality control measures were applied to the data. Variables (“features”) and subjects that had excessive missingness (i.e. more than 30% of variables) were excluded from clustering. Also excluded were features pertaining to our outcomes of interest; namely, incidence of parent-reported wheeze, asthma diagnosis and atopic disease at all timepoints. Feature selection was otherwise exploratory and all-inclusive, in that we attempted to retain as many individuals and variables as possible. We also included frequency of wheeze in the context of respiratory infection as it represented infection severity. This left us with a “complete-case” dataset of 186 subjects and 174 variables for clustering, which essentially covered variables from the first three years of life for each child. Some highly-skewed features, such as antibody and cytokine levels, were then subjected to logarithmic (base 10) transformation. Positional standardisation scaling was then applied across all variables. The complete list of clustering features is provided in **Supplementary Table 1**.

Unsupervised cluster analysis was performed on processed and scaled data using non-parametric expectation-maximisation (EM) mixture modelling (npEM) from the “mixtools” R package [40]. This method assumes that the frequency distributions of each cluster can be represented by non-parametric density estimates that are learned from the data in an iterative process. The optimal number of clusters was determined by scree plot and calculation of the Bayesian information criterion (BIC). The density functions generated by the resulting npEM model were then used to classify as many of the remaining “low-missingness” subjects as possible (31 of 36), so that the resultant groupings are a composite of unsupervised cluster analysis and supervised classification. Subjects were assigned to the cluster with ≥ 90% probability according to the model (**Supplementary Methods**).

Decision tree analysis was also conducted with the CAS clusters using “rpart” [41] to create classification trees that summarise inter-cluster differences and generate thresholds. We also specifically compared the classification trees with existing thresholds for atopy (any specific IgE at age 2 ≥ 0.35 kU/L, and/or any specific SPT at age 2 ≥ 2mm) [11], in terms of efficacy in predicting age-five wheeze.

The npEM clusters were then described and validated in two external datasets, MAAS (N=1085) [26] and COAST (N=289) [27]. This replication was performed by applying the density function-derived classification method used previously for the low-missingness CAS subjects. Only features that were common to both CAS and the replication cohorts (MAAS, COAST) were used for replicating the classification (**Supplementary Table 1**); these modified classification models were also tested in CAS, and the resulting CAS clusters compared to the pre-existing clusters in CAS. Further details can be found in the **Supplementary Methods** and **Supplementary Results**.

### 4.4 Statistical analyses

We performed statistical analyses comparing clusters in terms of all variables in the dataset. Of interest to us were our primary outcomes: asthma diagnosis and parent-reported wheeze at each timepoint. Comparisons were performed separately for each variable and timepoint. Statistical tests used included t-tests, Mann-Whitney-Wilcoxon tests, ANOVAs, Kruskal-Wallis tests, chi-squared and Fisher exact tests; and logistic and linear regression. For summary statistics, multiple testing adjustment was performed using the Benjamini-Yekutieli (BY) method, for all across-cluster tests (Cluster × trait); and for all comparisons between clusters (CAS1 vs. 2, 1 vs. 3, and 2 vs. 3). The BY method was chosen as it accounted for positive dependency across the highly-correlated variables in the CAS dataset [42]. For variables that underwent logarithmic transformation for statistical analyses before being transformed back, we used geometric means instead of arithmetic means to describe the measure of central tendency (in this case, the geometric mean is equivalent to the exponent of the arithmetic mean of the log-transform).

We then determined the predictors for age-five wheeze within each cluster. Both simple and multiple regression models were constructed; the former were built with and without a base set of covariates (sex, family history of asthma, BMI where available). The latter were built by manually selecting variables found to be most statistically-significant (at least p<0.05) in the univariate analyses, for each timepoint, followed by step-wise backward elimination to achieve the most parsimonious model with all predictors statistically-significant (p<0.05). Repeated-measures ANOVAs were also performed for selected predictors of age-five wheeze. Finally, generalised linear models (GLMs) were generated and their likelihood ratios examined using the “lrtest” function from the R package “Epidisplay” [43], to check how much cluster membership or classification trees improved upon prediction of age-five wheeze using selected predictors.

